# Circadian ribosome profiling reveals a role for the *Period2* upstream open reading frame in sleep

**DOI:** 10.1101/2022.08.09.503391

**Authors:** Arthur Millius, Rikuhiro Yamada, Hiroshi Fujishima, Kazuhiko Maeda, Daron M. Standley, Kenta Sumiyama, Dimitri Perrin, Hiroki R. Ueda

## Abstract

Many mammalian proteins have circadian cycles of production and degradation, and many of these rhythms are altered post-transcriptionally. We used ribosome profiling to examine post-transcriptional control of circadian rhythms by quantifying RNA translation in the liver over a 24-h period from circadian-entrained mice transferred to constant darkness conditions and by comparing ribosome binding levels to protein levels for 16 circadian proteins. We observed large differences in ribosome binding levels compared to protein levels, and we observed delays between peak ribosome binding and peak protein abundance. We found extensive binding of ribosomes to upstream open reading frames (uORFs) in circadian mRNAs, including the core clock gene *Period2 (Per2)*. An increase in the number of uORFs in the 5’UTR was associated with a decrease in ribosome binding in the main coding sequence and a reduction in expression of synthetic reporter constructs. Mutation of the *Per2* uORF increased luciferase and fluorescence reporter expression in 3T3 cells and increased luciferase expression in PER2:LUC MEF cells. Mutation of the *Per2* uORF in mice increased *Per2* mRNA expression, enhanced ribosome binding on *Per2*, and reduced total sleep time compared to that in wild-type mice. These results suggest that uORFs affect mRNA post-transcriptionally, which can impact physiological rhythms and sleep.

**Significance Statement:** *Period (Per)* is an iconic gene in the field of circadian rhythms since its discovery in 1971 by Seymour Benzer and Ronald Konopka in fruit flies. The inhibitory feedback loop of PER protein drives circadian rhythms. We show that *Per2* is regulated by an upstream open reading frame (uORF) in the 5’ untranslated region of *Period2* mRNA. Mutation of the *Per2* uORF altered the amplitude of luciferase reporter expression in well-characterized cell culture models. *Per2* uORF mutant mice had significantly elevated *Per2* mRNA levels and exhibited sleep loss, particularly during light-to-dark and dark-to-light transitions, which suggests a role for uORFs in modulating molecular and physiological circadian rhythms.

## Introduction

Life is remarkably adapted to the 24-hour rotational movement of the earth. In mammals, the molecular time-keeping mechanism for circadian rhythms relies primarily on a hierarchical network of transcription activators and repressors in cells and tissues (1). In the past, circadian clocks have been measured using systems approaches to quantify genome-wide changes in RNA levels (2), which has resulted in an understanding of the transcriptional regulatory network; however, less is known about how translation and post-transcriptional regulation influence biological rhythms.

Although 10% of genes are rhythmic in the liver (3), de novo transcription is responsible for only some this rhythmicity (4). For example, the timing of when a circadian mRNA is expressed does not necessarily correspond to that of mRNA translation or peak protein abundance (5). Some proteins have 24-h rhythms in abundance in the absence of rhythmic RNA expression (6, 7), which may suggest a role for rhythmic translation in regulating the clock (8).

In mouse liver, detection of low-abundant components of the core circadian circuit using systems proteomics is difficult (8, 9), unless special care is taken to examine a particular protein on a case-by-case basis (10) or by using advanced mass-spectrometry techniques (11, 12). Researchers have used next-generation sequencing of ribosome-bound mRNA protected from RNAse degradation to understand how translation regulation affects protein output (13, 14). Previous studies using ribosome profiling to measure daily rhythms focused on a cell culture model (15) or mouse tissues in light-dark conditions (16–18) to examine rhythms in diurnal gene expression, which may be influenced by non-circadian time-keeping systems. These studies also examined the timing between RNA abundance and ribosome binding, but it remains unclear how circadian translation relates to peak protein abundance in terms of protein turnover and timing. For example, the peak abundance of PER and CRY proteins are delayed relative to expression of their mRNA in the liver (19) and suprachiasmatic nuclei (SCN) (20). This difference in timing may result from a delay in RNA processing before translation, a delay during translation, a delay in protein transport from one cellular location to another, or a delay in protein turnover (21, 22).

Previously, we developed a mass spectrometry method called MS-based Quantification By isotope-labeled Cell-free products (MS-QBiC) to determine the absolute protein levels of 16 selected circadian proteins in mice liver over a 24-h period (10). This method takes advantage of the PURE system (23) for reconstituting cell-free protein expression of optimal peptide standards for detection and quantification by selected reaction monitoring (SRM)-based targeted proteomics analysis. We found delays between the peak levels of RNA expression, as measured by quantitative PCR (qPCR), and the abundance of the corresponding protein, suggesting either a delay in post-transcriptional RNA processing or in protein turnover. Here, we investigated the same liver samples by ribosome profiling in order to understand the timing of ribosome binding compared to peak protein and RNA levels. We found that upstream open reading frames (uORFs) modulated translation globally, repressed reporter expression in a combinatorial manner, and suppressed expression of *Per2*. Mutation of the *Per2* uORF in mice reduced total sleep duration, particularly during the early morning and early evening, without disrupting the circadian period, which suggests that uORF-mediated
repression may impact physiological behaviors.

## Results

### Ribosome profiling of liver from mice in constant darkness conditions

An experimental workflow was designed to analyze ribosome-protected mRNA fragments from liver samples previously examined by MS-QBiC (10) (*SI Appendix*, Fig. S1). Briefly, mice were entrained to a 12-h light/12-h dark (LD) cycle for 14 days, transferred to constant darkness (DD) for 24 h, and sacrificed at circadian times (CT0, CT4, CT8, CT12, CT16, CT20, and CT24). Liver samples from two mice were collected and analyzed following established ribosome profiling protocols (24). We prepared ribosome profiling libraries and sequenced ∼70 million reads per sample; this resulted in 25-45 million reads that could be mapped to mRNA (*SI Appendix*, Table S1). Ribosome-protected fragments primarily aligned to the coding regions and 5’ untranslated regions (5’UTRs) of the mRNA, with few reads mapping to the 3’ untranslated region (3’UTR) (*SI Appendix*, Fig. S2A). Alignment of coding sequence (CDS)-mapped reads based on the footprint length revealed reading frame periodicity (*SI Appendix*, Fig. S2B). Reads were of the expected size, mapped with a high percentage to mRNA, and were correlated between samples (*SI Appendix*, Fig. S2C–E).

From ∼14,000 well-translated transcripts (defined by a median of at least two reads per five-codon mRNA window), we identified rhythms in ribosome-protected read fragments using the JTK_CYCLE algorithm (25), yielding 2952 rhythmic transcripts with an adjusted P < 0.05, including well-known circadian transcripts such as *Bmal1, Per1, Per2, Clock*, and *Cry1* (*SI Appendix*, Dataset S1, Table S2, Fig S6A).

### Relationship between ribosome profiling reads and protein levels

We compared the timing of protein production, as measured by ribosome profiling reads, to the absolute number of protein molecules per cell for 16 previously reported core circadian proteins (10). There was broad agreement in the timing of ribosome binding compared to that of protein abundance (Fig. 1A). However, for several circadian proteins, such as BMAL1 and CLOCK, there was a delay of approximately 6 hours between peak ribosome binding and peak protein abundance, clearly outside the 4-h range of our sampling intervals (Fig. 1B), which suggested that there is a post-translational delay in protein turnover for these molecules rather than a post-transcriptional delay in ribosome binding. We compared the average number of ribosome profiling reads to the number of protein molecules over a 24-h period, but found that the two values were poorly correlated (Fig. 1C). We measured this protein production efficiency (as defined by the ratio of our mass-spectrometry protein levels to ribosome profiling reads) at each time point. For some proteins, such as BHLHE40, a large amount of ribosome binding resulted in a moderate amount of protein (Fig. 1D); whereas for other proteins, such as PER2, a much smaller amount of ribosome binding resulted in the same amount of protein as BHLHE40 (Fig. 1E), which suggests that BHLHE40 protein is more unstable than PER2 or that some other factor is limiting the amount of BHLHE40 produced. Thus, ribosome profiling reads can provide an approximate estimate for when a protein is produced, but protein abundance reflects both protein production and post-translational mechanisms to control overall protein levels (26).

**Figure 1.**
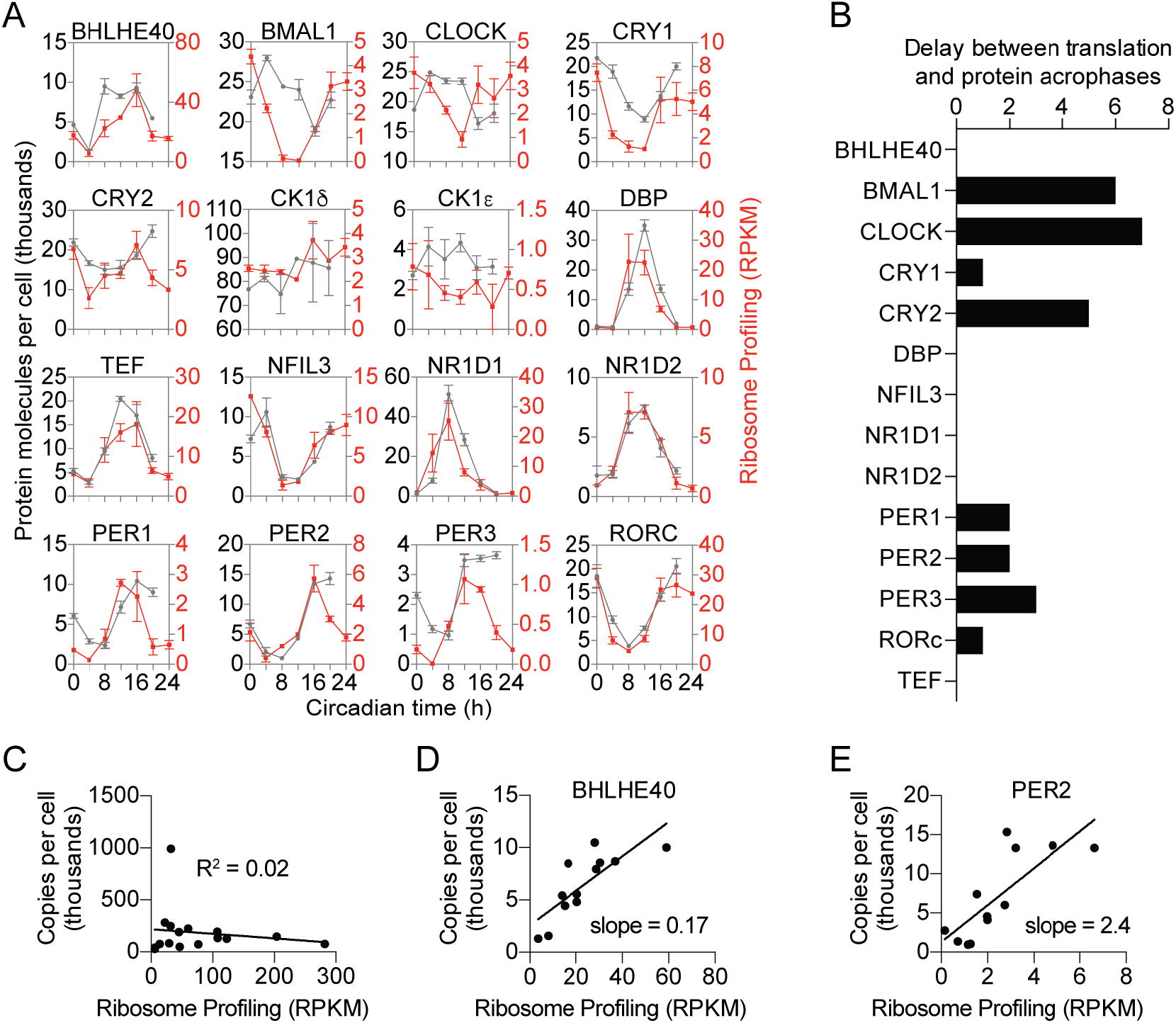
Measuring translation and protein from the same liver samples for 16 circadian mRNAs. (A) Time course of the quantified proteins and the corresponding translation level. Protein concentration (gray) is expressed as copies per cell assuming the total amount of protein per cell is 0.5 ng for the average for all quantified peptides (10). Ribosome binding (red) is expressed as reads per kilobase of transcript per million mapped reads (RPKM). Data are mean ± SEM. (B) Phase delay between ribosome binding and protein abundance. Copies per cell were averaged from all peptides for each protein. JTK analysis was used to estimate the phases for these averaged protein levels and for ribosome binding RPKM levels. (C) Linear regression analysis comparing ribosome binding to protein amount averaged over 24 hours showed that there was no correlation between ribosome binding and protein abundance (D and E) Linear regression analysis comparing ribosome binding to protein amount at each sampled time point showed that some proteins like BHLHE40 (D) had high protein turnover or reduced protein biosynthesis whereas other proteins like PER2 (E) had low protein turnover or increased protein biosynthesis.

When we compared mRNA and protein abundance phases in our dataset in DD to the reported phases in previously published studies of mouse liver in LD (12, 15, 16, 18), we observed a correlation with a 1-2 h delay in DD (*SI Appendix*, Fig. S10A, C). The transcripts with the largest differences in delay in the mRNA-to-peak-protein abundance between LD and DD conditions included *Nfil3, Per2, Per3*, and *Bmal1* (*SI Appendix*, Fig. S10B). The differences resulted from a slightly earlier peak in RNA abundance and ribosome binding in LD rather than a delay in the protein abundance peak; there was little difference between LD and DD in the delay between RNA abundance and ribosome binding (*SI Appendix*, Fig. S10D). Although these small differences may reflect a true biological effect from light-dependent changes in transcription, they could also simply result from differences in the algorithms used for phase determination, differences in experimental conditions, or both. Besides these 16 core circadian transcripts, most other transcripts with rhythmic ribosome binding in our dataset had a phase of around CT0 (*SI Appendix*, Fig. S6).

### Upstream open reading frames (uORFs) suppress translation

We also noticed a relationship between ribosome occupancy and the presence of upstream open reading frames (uORFs), similar to previous reports (15, 18). Roughly half of all mouse transcripts contained at least one uORF (*SI Appendix*, Fig. S4B), and we found rhythmic ribosome binding in 602 uORFs with an adjusted P < 0.05 using JTK_cycle (*SI Appendix*, Dataset S2). In particular, for circadian transcripts, such as *Cry1* and *Bmal1*, there appeared to be increased ribosome binding in uORF regions, as measured by increased ribosome profiling reads (Fig. 2A), although ribosome binding in the 5’UTR was not always associated with uORFs in circadian transcripts (*SI Appendix*, Fig. S4A). mRNAs with increased numbers of uORFs had lower levels of ribosome occupancy in the downstream CDS (Fig. 2B), whereas the length of the uORF and the distance of the uORF to the start codon did not have a significant impact on ribosome binding in the CDS (*SI Appendix*, Fig. S4C, D). To investigate whether uORFs were sufficient to suppress translation in a combinatorial manner, we created a luciferase reporter vector with multiple synthetic uORFs. Predictably, increasing the number of uORFs reduced luminescence from the reporter (Fig. 2C). Using this synthetic reporter, we varied the position of a single uORF relative to the start codon or the uORF length and found a higher degree of uORF-mediated repression the closer the uORF was to the start codon; however, uORF length had no effect on repression (*SI Appendix*, Fig. S4E, F). Both the position of the uORF and the number of uORFs altered the amplitude and mesor (mean luminescence) without altering the period using two different promoters in cell-based circadian luminescence assays (*SI Appendix*, Fig. S15). Other factors, such as the Kozak consensus of the uORF, overlap with the CDS, or uORF conservation may also impact repression strength (27, 28).

**Figure 2.**
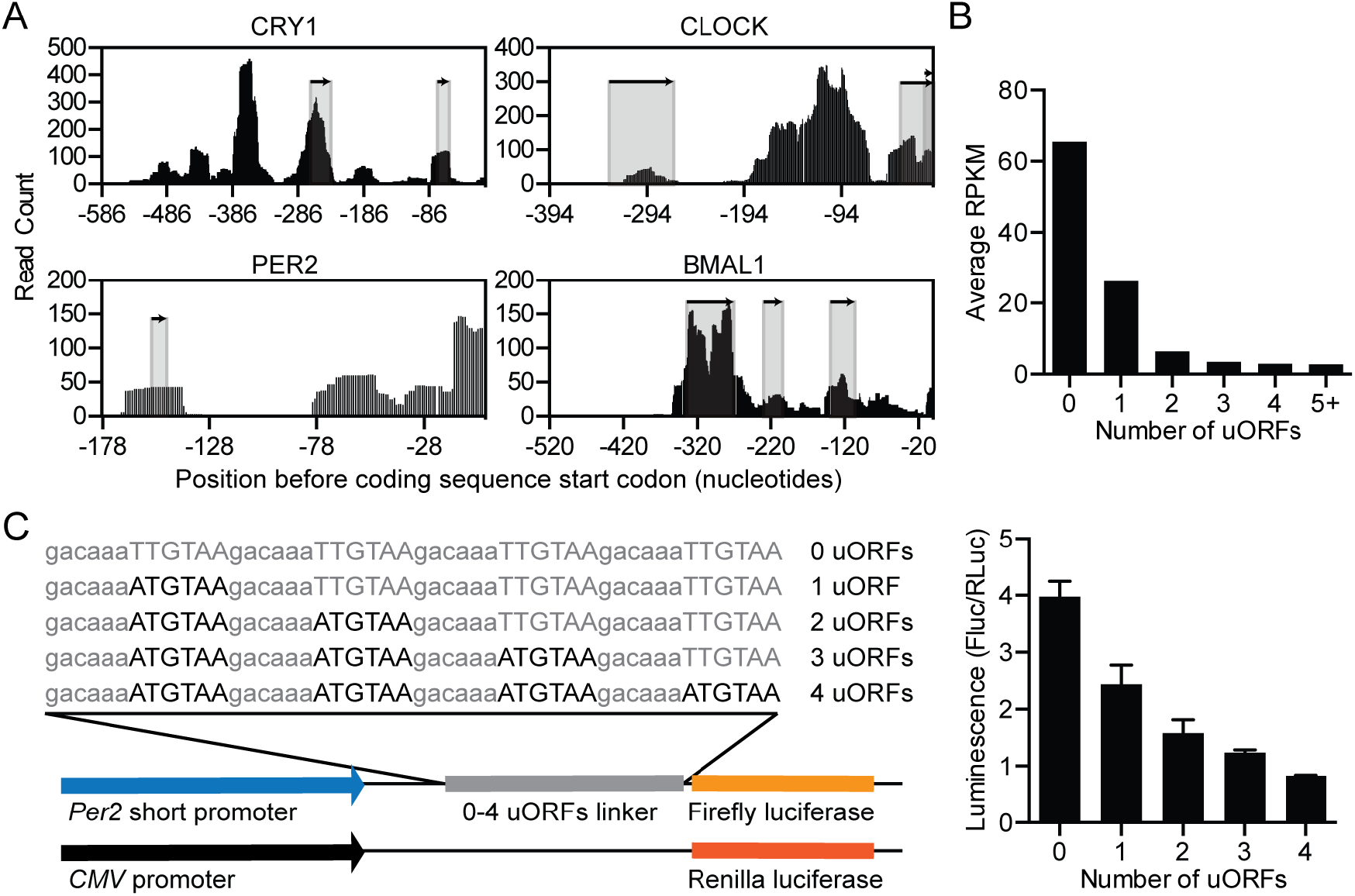
Upstream open reading frames (uORFs) suppress protein expression. (A) The 5’UTRs of *Cry1, Clock, Per2*, and *Bmal1* transcripts contain at least one uORF (shaded regions). Distribution of raw read counts from ribosome profiling data (black bars) show ribosomes bounds in regions corresponding to uORFs. (B) Global distribution of uORFs compared to ribosome profiling translation level (RPKM) shows that mRNAs with more uORFs are translated less compared to mRNAs with fewer uORFs. (C) Introduction of a variable number of synthetic uORFs represses relative luminescence from a *Per2* short promoter in a dose-dependent manner.

### The Per2 uORF suppresses reporter expression without altering period

To further explore the impact of uORFs on circadian rhythms, we focused on the uORF in the circadian transcript *Per2*. This uORF is evolutionarily conserved and consists of only a start and stop codon (Fig. 3A and *SI Appendix*, Fig. S13), which eliminates any potential effect of a translated uORF peptide on the regulation of *Per2*. Ribosomes bound the *Per2* uORF rhythmically and slightly before peak ribosome binding on the *Per2* transcript (Fig. 3B). Mutation of the uORF in *Per2* increased the amplitude of expression without affecting the phase or period in 3T3 cells transfected with a luminescence reporter (Fig. 3C, D). This increase in amplitude was not affected by the amount of transfected plasmid, inclusion of the full-length *Per2* 5’UTR, or addition of PER2 protein (*SI Appendix*, Fig. S5). CRISPR/Cas9-mediated mutation of the *Per2* uORF in PER::LUC MEFs increased overall luciferase expression levels compared to that of wild-type MEFs; however, we were unable to generate circadian rhythms in these cells to determine the impact of the *Per2* uORF on other circadian parameters such as period and phase (*SI Appendix*, Fig. S12).

**Figure 3.**
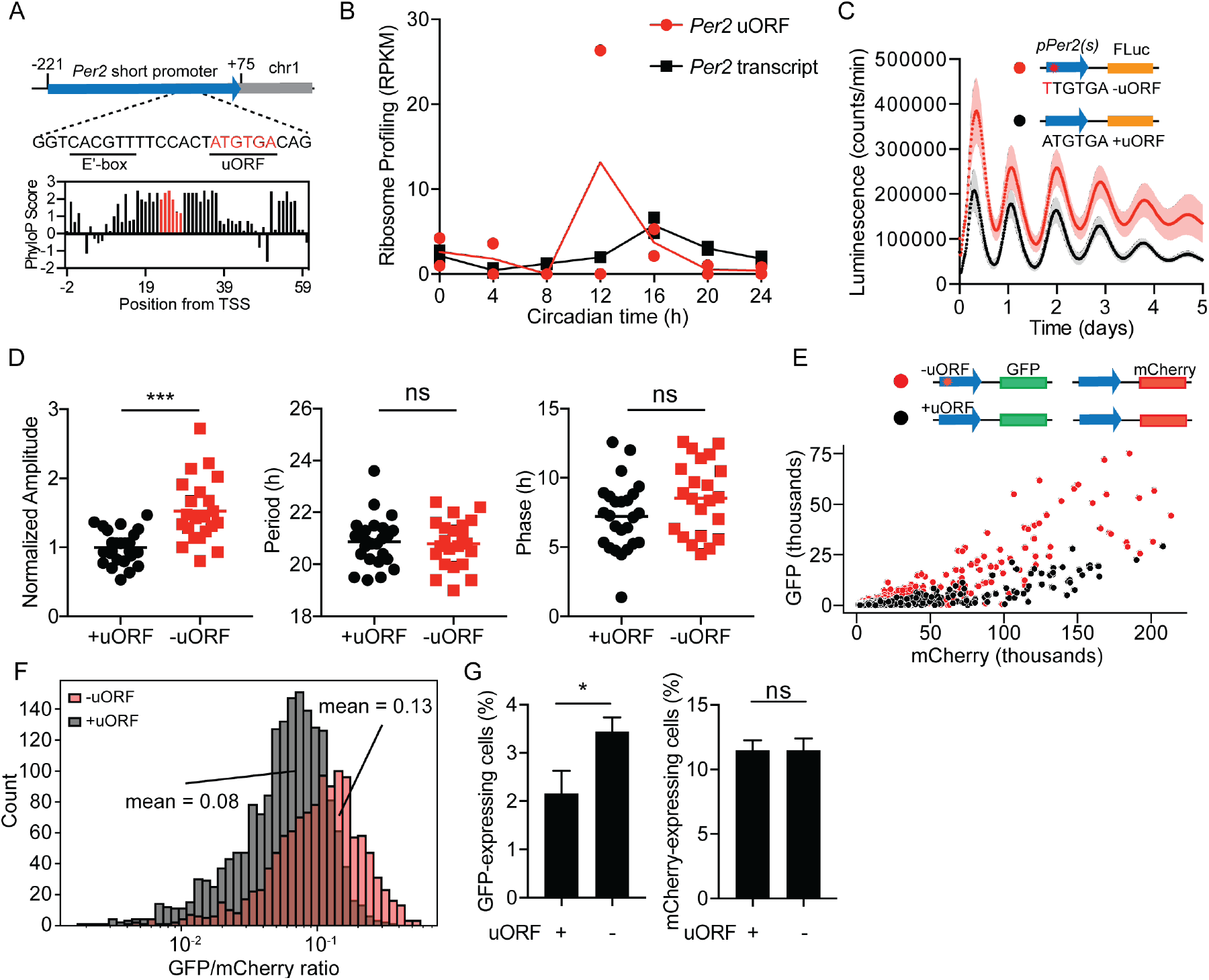
The *Per2* uORF is evolutionarily conserved and suppresses PER2 expression. (A) Genomic position of the Per2 short promoter, which contains an E’box and uORF. Evolutionary conservation scores according to UCSC genome browser among 22 mammalian species in the *Per2* 5’UTR (black bars) with the uORF highlighted (red bars). (B) Ribosome binding (RPKM) on the *Per2* uORF (red) compared to the *Per2* transcript (5’UTR, CDS, and 3’UTR, black). (C) Representative bioluminescent traces showing that mutation of the *Per2* uORF (ATGTAA to TTGTGA) increases the expression level of a pGL3-P(*Per2*)-d*Luc* luminescent reporter. Reporter containing uORF (black), reporter with a mutant uORF (red). Shaded region is SD. (D) Cosinor analysis showing the normalized amplitude (*left*), period (*middle*), and phase (*right*) for cells transfected with the *Per2* uORF (black) or a mutant *Per2* uORF (red). Data from each group comprise at least 22 traces from 8 different experiments (see *SI Appendix*, Fig. S5 for the plots of individual traces for each experiment). (E) Representative scatterplot of GFP versus mCherry expression for 3T3 cells transfected with a GFP-fluorescence reporter with a *Per2* uORF (+uORF, black) or a mutant *Per2* uORF (-uORF, red) and a mCherry-normalization plasmid. The analyzed cells were gated on mCherry+ expression. (F) Histogram of the GFP/mCherry ratio for cells in (E). (G) The percentage of GFP+ cells (*left*) was significantly higher in the -uORF population compared to the +uORF population, whereas the percentage of mCherry+ cells (*right*) was unchanged (n = 3 independent experiments). See *SI Appendix*, Fig. S11 for FACS gating strategy and analysis of cells from the GFP+ gate.

To understand uORF-mediated repression in individual cells, we created a GFP fluorescent reporter plasmid driven by the *Per2* promoter with or without a mutation in the *Per2* uORF. We transfected 3T3 cells with these plasmids and an mCherry normalization plasmid and analyzed the cells by FACS (*SI Appendix*, Fig. S11). Mutation of the uORF increased GFP brightness without affecting mCherry brightness in individual cells (Fig. 3E), and increased the GFP-to-mCherry ratio of the population (Fig. 3F) regardless of whether the cells were first gated on mCherry expression (Fig. 3E, F) or GFP expression (*SI Appendix*, Fig. S11C, D). Mutation of the uORF also increased the total number of GFP-expressing cells without affecting the total number of mCherry-expressing cells (Fig. 3G). Thus, the *Per2* uORF can repress reporter expression within individual cells to control circadian amplitude.

### Mice with a mutation in the Per2 uORF have reduced sleep

We used CRISPR/Cas9 to generate a knock-in mouse harboring a mutation in the uORF of *Per2*, which removed both the start and stop codon of the uORF without disrupting a nearby E’-box (*SI Appendix*, Fig. S7A). Wild-type and mutant mice were phenotyped over 13 days in LD and then for 12 days in DD using the Snappy Sleep Stager (29, 30), which is a respiration-based method to characterize sleep/wake parameters and circadian rhythms. Both male and female *Per2* uORF mutant mice had significantly reduced sleep per day (mean ± SEM: 717 ± 19 min and 642 ± 7 min, respectively) compared to their wild-type littermates (774 ± 11 min and 678 ± 4 min, respectively, P < 0.001 by two-way ANOVA), and a corresponding increase in wake duration per day (Fig. 4A, B and *SI Appendix*, Fig. S8A) in LD conditions. There was a decrease in sleep episode duration and a significant increase in the transition probability from sleep to awake termed *P*_*sw*_ in *Per2* uORF mutant mice (*SI Appendix*, Fig. S8B); however, there were no differences in other sleep parameters such as amplitude, *P*_*ws*_ (the transition probability from awake to sleep), or wake episode duration (*SI Appendix*, Fig. S8C). *Per2* uORF mutant mice exhibited reduced sleep duration particularly during the light-to-dark and dark-to-light transitions in the early morning and early evening (Fig. 4C, D). There were also differences in sleep duration later in the morning (*SI Appendix*, Fig. S8D), but not at other times during the day (*SI Appendix*, Fig. S8E). Next, we observed mice in constant darkness over 12 days and found no differences in activity rhythms or period (Fig. 4E); furthermore, there were less pronounced sleep differences in DD compared to LD (*SI Appendix*, Fig. S9). Cosinor analysis of the activity rhythms in LD and DD conditions revealed no differences in period and amplitude between wild-type and *Per2* uORF mutant mice (*SI Appendix*, Fig. S9).

**Figure 4.**
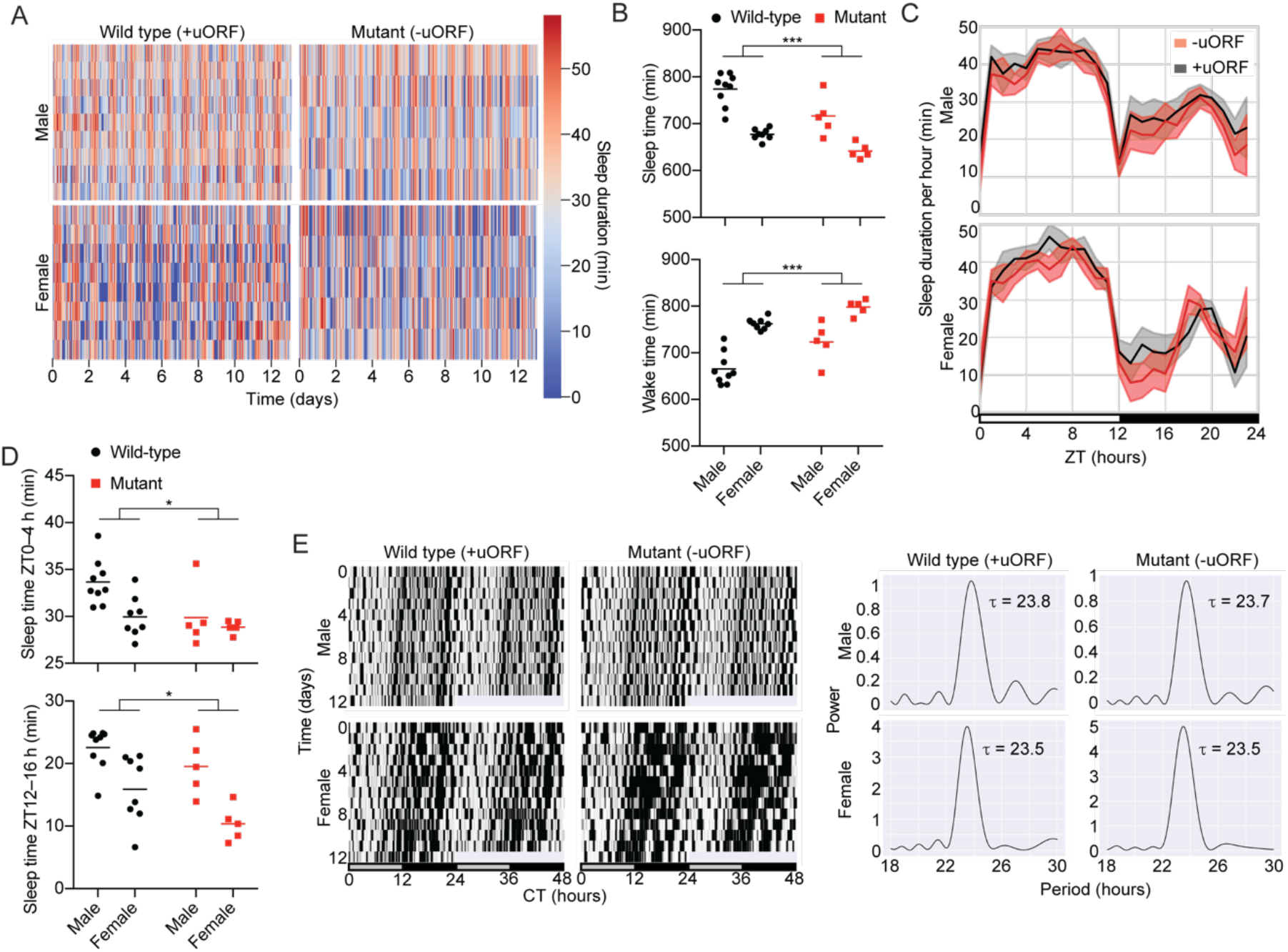
Disruption of the *Per2* uORF reduces sleep in mice. (A) Sleep duration per hour over 13 d in 12-h light/12-h dark (LD) conditions for *Per2* uORF mutant and wild-type male and female mice. Each row indicates data from one mouse. (B) Mean sleep (*top*) and wake (*bottom*) duration over 24 h, averaged over 13 d. Red, *Per2* uORF mutant mice. Black, wild-type. (C) Sleep duration per hour over 24 h in LD, averaged over 13 d for *Per2* uORF mutant (red) and wild-type (black) male (*top*) and female (*bottom*) mice. Lines indicate mean sleep duration at each time of day for each strain. Shaded area, SD at each time point. (D) Mean sleep duration per hour averaged over 13 d during the early morning (*top*) and early evening (*bottom*). Per2 uORF mutant (red) and wild-type (black) dots are individual mice; lines indicate mean. (E) Representative double-plotted actograms of sleep duration per 6 min bins of wild-type and *Per2* uORF mutant mice in constant darkness (DD) over 12 d (*left*) and corresponding periodograms (*right*).

To understand the molecular mechanisms behind these phenotypic differences, we performed ribosome profiling and total RNA sequencing of liver from wild-type and *Per2* uORF mutant male mice sacrificed at ZT02-04 (*SI Appendix*, Fig. S18, 19, Dataset S4, S5). Because *Per2* levels are near their lowest at ZT02-04, we hypothesized that this time point would show the largest difference in derepression of *Per2* by mutation of the *Per2* uORF. There was an increase in ribosome binding in *Per2* uORF mutant mice (*SI Appendix*, Fig. S19C), particularly in the 5’UTR and around the start codon (*SI Appendix*, Fig. S19D, E). *Per2* RNA levels were also higher in the mutant mice by total RNA sequencing (*SI Appendix*, Fig. S19C, H and Dataset S5) and confirmed by qPCR analysis (*SI Appendix*, Fig. S19B), although no differences were observed in *Per2* translation efficiency (TE), which is defined as the ratio of ribosome profiling reads to bulk RNA-seq reads (*SI Appendix*, Fig. S19G and Dataset S4). Protein levels were similar in *Per2* uORF mutant mice compared to that in wild-type mice (*SI Appendix*, Fig. S19A, I). Taken together, these results suggest that *Per2* mRNA expression and ribosome binding is suppressed by the *Per2* uORF, which has a moderate impact on sleep duration.

## Discussion

Global proteomics studies of mice livers revealed large numbers of diurnally rhythmic proteins. It has been estimated that 20–50% of these proteins are dependent on post-transcriptional and post-translational mechanisms (8, 9, 11, 12, 31). Previously, we used targeted proteomics to detect the rhythmicity and concentration of a select number of low abundance core circadian proteins in constant darkness conditions (10). Here, we used ribosome profiling and quantitative mass spectrometry to understand the relationship between those circadian protein rhythms and mRNA translation.

We found a correlation between the timing of protein production, as measured by ribosome profiling, and peak protein abundance for most transcripts (Fig. 1). However, BMAL1 and CLOCK proteins had a delay between peak ribosome binding and protein levels (Fig. 1) caused by delayed protein turnover via post-translational modification pathways (32), rather than a delay in ribosome binding (*SI Appendix*, Fig. S3). The peak protein abundance of PER and CRY is also slightly delayed relative to their peak mRNA abundance (10, 19, 20), and we found this was also due to protein turnover rather than to a delay in ribosome binding (*SI Appendix*, Fig. S3), similar to previous ribosome profiling studies (15, 16, 18). There was a larger delay between the peak abundance of *Cry2* ribosome binding and CRY2 protein than that of *Cry1*, which may simply be a result of the lower amplitude of CRY2 rhythms.

We lacked the RNA-seq data needed to compute TE in our circadian samples, which limited our ability to determine how uORF features in 5’UTRs, such as number, length, and distance to CDS, alter translation efficiency. However, using the closest publicly available RNA-seq data (4) to our ribosome profiling data as a proxy for transcript abundance (*SI Appendix*, Dataset S3), we observed a decrease in TE in transcripts with one or more uORFs (*SI Appendix*, Fig. S16), but no relationship between TE and uORF length or uORF distance to CDS (*SI Appendix*, Fig. S17).

Most transcripts with rhythmic translation in our dataset had a phase around CT0, corresponding to what would be the dark-to-light transition (*SI Appendix*, Fig. S6). Less than 10% of the circadian translatome had significant rhythms in mRNA abundance according to JTK cycle analysis (P < 0.05), and these transcripts were enriched for circadian and metabolic processes by Panther gene ontology (GO) analysis (Fig. S6B, C). Transcripts with a non-significant mRNA rhythmicity by JTK cycle analysis (P > 0.05) also were biased toward CT0, but there was no clear GO biological process underlying these rhythms (*SI Appendix*, Fig. S6D, E). The set of transcripts with phases between CT22 and CT02 (*SI Appendix*, Fig. S6F, G) included *Npas2, Bmal1, Cry1, Nfil3*, and *Clock* are all regulated by the nuclear receptor NR1D1, also known as Rev-erbα, which transcriptionally represses many genes involved in metabolism in a tissue- and circadian-dependent manner (33). In the liver, NR1D1 predominantly binds genomic locations at ZT08-ZT10 with little-to-no binding at ZT22 (34–36). Thus, NR1D1 derepression may be driving some of transcriptional, and subsequent translational, peak observed in our data between CT22 and CT02. In addition, cyclic changes in translation machinery and polyadenylation (37, 38), translation activity of BMAL1 mediated by the mTORC1/S6K1 (39), or mTORC1 regulation by PER2 (40) may also drive rhythmic translation of non-rhythmic mRNA transcripts.

We observed widespread binding of ribosomes to uORFs in the 5’UTRs of many core circadian transcripts and found an inverse relationship between uORF number and ribosome occupancy or luciferase expression in transiently transfected reporters or CRISPR/Cas9-generated MEF cells (Fig. 2, *SI Appendix*, Fig. S4, Fig. S12, and Fig. S15). Approximately 50% of mouse and human mRNAs contain uORFs, which are associated with widespread translational repression (28). Thus, uORF-mediated translation suppression may constitutively suppress the abundance of circadian proteins or provide a post-transcriptional foothold to adjust protein abundance by altering the activities of ribosome reinitiation factors (41–43). Moreover, mutagenesis of uORFs have been shown to alter rhythmicity in other circadian systems (44, 45). Not all ribosome binding in 5’UTRs was associated with uORFs, and we observed extensive ribosome binding in areas without an apparent uORF for *Cry1, Nr1d1, Clock, Dbp*, and *Per3* (Fig. 2 and *SI Appendix*, Fig. S4). For the *Nr1d1* 5’UTR, we observed close overlap in the nucleotides necessary for internal ribosome entry site (IRES)-mediated translation (46) and ribosome binding (*SI Appendix*, Fig. S14A), and confirmed IRES-mediated translation from the *Nr1d1* but not the *Cry1* 5’UTR (*SI Appendix*, Fig. S14B). Near-cognate uORF translation (from a non-AUG start codon) may also drive ribosome binding (14). In silico mapping of near-cognate start codons with ribosome binding data revealed numerous potential near-cognate uORFs (*SI Appendix*, Fig. S14C) but experimental validation is still needed.

The uORF in the *Per2* 5’UTR is an attractive target to understand the role of uORFs in circadian biology, not only because cis-elements within the *Per2* promoter are well-understood (47–50), but also because the *Per2* uORF is too short to encode a peptide. PER2 production is post-transcriptionally controlled by miRNAs (51, 52), antisense transcription (53), and hnRNP1-mediated mRNA degradation (54), which suggests that *Per2* post-transcriptional control is important for producing the optimal amount of PER2 protein. Mutation of the *Per2* uORF increased the amplitude of a reporter plasmid without affecting the phase or period (Fig. 3), similar to effects observed by mutating the *Per2* antisense transcript (55).

Both the abundance and timing of *Per2* expression are critical for maintaining circadian rhythmicity in mice because constitutive expression of PER2, unlike CRY1, disrupts behavioral rhythms (56). Several post-transcriptional mechanisms such as *Per2* anti-sense transcription (53) and miRNAs (51, 52, 57) in the *Per2* genomic locus alter the timing and abundance of PER2 protein expression. For example, replacing the *Per2* 3’UTR with a SV40 late poly(A) signal greatly amplifies bioluminescence rhythms in PER2:LUC mice and increases free-running periods (52). Mutation of the *Per2* uORF increased amplitude in reporter cells (Fig. 3), but did not increase the free-running period in mice (Fig. 4). *Per2* uORF mutant mice did have reduced total sleep in LD conditions (Fig. 4), as observed in *Per1/Per2* double mutant mice under similar sleep phenotyping conditions (29), but no significant change in sleep in DD conditions (*SI Appendix*, Fig. S9). Although our *Per2* mutation abolished the canonical ATGTGA uORF in the *Per2* 5’UTR, it also introduced a near-cognate uORF (CTGTAG) two nucleotides upstream of the original uORF. We think this near-cognate uORF was non-functional because we observed increased *Per2* ribosome binding and mRNA levels in *Per2* uORF mutant mice compared to that in wild-type mice (*SI Appendix*, Fig. S19) and increased luciferase expression in mutant PER2:LUC MEF cells (*SI Appendix*, Fig. S12). One possible explanation for the rise in *Per2* mRNA levels is uORF-triggered nonsense mediated decay (NMD) (58). In this scenario, mutation of the *Per2* uORF derepresses NMD of *Per2*, which results in an increase in *Per2* mRNA. However, *Per2* mRNA expression does not increase upon knockdown of the NMD component *Smg6* (59), which suggests that other proteins in NMD independent of SMG6 may be involved (60). Another group deleted the start codon of the *Per2* uORF, but found no effect on *Per2* mRNA levels and did not perform ribosome profiling (61), so it is unclear if ribosome binding in *Per2* is elevated in their *Per2* uORF mutant mice as it is in our mutant mice. In both studies, PER2 protein levels in the mutant mice were similar to that in wild-type mice, indicating that circadian proteostasis pathways compensate for the increase in PER2 protein production. In LD conditions, sleep in our *Per2* uORF mutant mice was particularly reduced during light-to-dark and dark-to-light transitions (Fig. 4), which may result from a lower sleep episode duration and an increase transition probability from sleep to wake (*SI Appendix*, Fig. S8B). Regulation of PER2 stability affects sleep (62, 63), and sleep deprivation reciprocally affects *Per2* expression (64–67) providing a rationale for how increased *Per2* expression could disrupt sleep in our mice. Mutation of the E’box cis-element, which is only a few base pairs upstream of the *Per2* uORF, abolishes molecular oscillations in mutant cells without disrupting the free running period in mice (47). These mice re-entrain quicker under an artificial jetlag experiment than wild-type mice, and it will be interesting to observe how *Per2* uORF mutant mice behave under similar conditions. Further studies, particularly at a neurological level, are also needed to understand the precise mechanism by which sleep is reduced in these animals.

## Materials and Methods

### Methods

#### Animals

All animal experiments were approved by the Institutional Animal Care and Use Committee of the RIKEN Kobe branch. Eight-to-ten week-old wild-type male mice (C57BL/6N, Japan SLC) were entrained under 12-h light (400 lx) 12-h dark (LD) for two weeks. Twenty-four hours after transferring to constant darkness (DD), wild-type mice were sacrificed every 4 h over one day (CT0, CT4, CT8, CT12, CT16, CT20, and CT24) for ribosome profiling analysis as in (10). Livers were excised, snap frozen in liquid nitrogen, and stored at -80 °C until use. For details regarding *Per2* uORF mutant mouse construction, see the *SI Appendix*.

#### Plasmids

For details regarding plasmid construction, see the *SI Appendix*.

#### Ribosome Profiling

Ribosome profiling was performed essentially as described in (24). Frozen liver samples (∼50 mg) were pulverized and then homogenized in 400 μl polysome lysis buffer (150 mM NaCl, 20 mM Tris-HCl pH 7.4, 5 mM MgCl_2_, 5 mM DTT, 100 μg/ml cycloheximide, 1% Triton X-100, 25 U/ml Turbo DNAse I). Lysates were incubated on ice for 5-10 min, triturated through a 26-G needle 10 times, clarified by centrifugation (20,000 × *g* for 10 min at 4 °C), and 300 μl of supernatant was transferred to a new tube. Unbound RNA was digested by the addition of 7.5 μl RNAse I (ThermoFisher) for 45 min and then stopped by 10 μl SUPERase In RNAse Inhibitor (ThermoFisher). The digestion was transferred to 13 mm × 51 mm polycarbonate ultracentrifuge tubes, layered on top of 0.9 ml sucrose cushion (150 mM NaCl, 20 mM Tris-HCl pH 7.4, 5 mM MgCl_2_, 5 mM DTT, 100 μg/ml cycloheximide, 1 M sucrose, 10 U/ml SUPERase In), centrifuged in a TLA100.3 rotor at 70,000 rpm at 4 °C for 4 h. Ribosome pellets were resuspended in 0.7 ml Qiazol and mRNA was recovered using the miRNeasy RNA extraction kit (Qiagen) according to the manufacturer’s instructions. For library construction and sequencing, see the *SI Appendix*.

#### Bioinformatic analysis of ribosome profiling

For details regarding sequence processing and alignment, see the *SI Appendix*. For each transcript, non-overlapping five-codon windows were tiled across the coding region, and the transcript was considered well-translated if it had a median value of at least 2 reads per window (excluding the excluding the first fifteen and the last five codons). Using all well-translated transcripts, we created file where each transcript is represented across all samples by its RPKM value. We used this file in JTK_CYCLE (25) to identify all rhythmic transcripts. We used a threshold on the adjusted P < 0.05 to assess significance.

To detect uORFs, we processed the 5’UTR of all transcripts, extracted their sequence, and identified all pairs of a start codon (AUG) and a stop codon (UGA, UAA, or UAG) in phase with each other. Using the same read assignment method as in the *SI Appendix*, we also allocated reads aligned to the UTR to their corresponding uORF (where appropriate).

#### Cells

NIH3T3 and MEFs from PER2::LUCIFERASE (PER2::LUC) knock-in reporter mice (68) were cultured in DMEM (ThermoFisher), 10% FBS (JRH Biosciences), and 1% penicillin/streptomycin (ThermoFisher) at 37 °C in 5% CO_2_. For additional details, see the *SI Appendix*.

#### Real-time Circadian Luciferase Assay

Real-time circadian luciferase assays were performed as previously described (69). Briefly, the day before transfection, NIH3T3 cells or PER2:LUC MEFs were plated onto 35-mm dishes at a density 4 × 10^5^ per well. The following day NIH3T3 cells were co-transfected using FuGene6 (Roche) with 1 μg of the indicated luciferase reporter plasmid according to the manufacturer’s instructions (n = 3) and cultured at 37 °C. After 72 h, the media in well was replaced with 2 ml of DMEM containing 10% FBS supplemented with 10 mM HEPES (pH 7.2, ThermoFisher), 0.1 mM luciferin (Promega), antibiotics, and 10 μM forskolin (Fermentek, NIH3T3) or 100 μM dexamethasone (ThermoFisher, PER2::LUC MEFs). Luminescence was measured by a photomultiplier tube (LM2400R, Hamamatsu Photonics) for 1 min at 12 min intervals in a darkroom at 30 °C.

#### Dual-Luciferase Assay

NIH3T3 cells were plated on 6-well plates at a density of 2 × 10^5^ per well. The following day cells were co-transfected with 0.95 μg of a Firefly *luciferase* reporter plasmid and 50 ng phRL-SV40 plasmid (Renilla luciferase, Promega) as an internal control for transfection efficiency using FuGene6 (Roche). Cells were harvested and assayed by the Dual-Luciferase Reporter Assay System (Promega) according to the manufacturer’s instructions 48 h after transfection.

#### Preparation of mouse nuclear lysate and immunoblot analysis

Mice were sacrificed by cervical dislocation, and livers were dissected at CT2-4, snap frozen in liquid nitrogen, and stored at -80 °C. Liver extracts were prepared according to (70) with minor modifications. See the *SI Appendix* for further details.

#### FACS

NIH3T3 cells were plated onto plastic 35-mm dishes or 35-mm imaging dishes (Ibidi) at a density 4 × 10^5^ per well. The following day cells were co-transfected using FuGene6 (Roche) with 0.5 μg of the indicated fluorescence reporter plasmids (1 μg total) according to the manufacturer’s instructions (n = 3) and cultured at 37 °C. After 72 h, cells were trypsinized, sorted using FACSAria I (BD Biosciences), and analyzed by FlowJo version 10.8.1.

#### Sleep Phenotyping

Sleep phenotyping was conducted in in 12-week-old (LD) and 14-week-old (DD) mice in a Snappy Sleep Stager (SSS) using *Per2* uORF mutant mice and wild-type littermates as a control (29). SSS is a non-invasive, respiration-based sleeping staging system in which mice are placed in a chamber connected to a respiration sensor that detects pressure differences between the outside and inside of the chamber. Detailed methods have been described previously (29). For details regarding sleep and wake parameters, see the *SI Appendix*.

#### Statistical Analyses

Statistical analyses were performed in R version 3.4.3, Prism 7.0, and custom Jupyter notebooks version 6.3. Two-way ANOVA and a Student’s t-test was used to test differences in sleep parameters between wild-type and *Per2* uORF mutant mice. For the Student’s t-test, data were confirmed gaussian-distributed by a Shapiro-Wilk normality test and an F test was used to check equality of variances. Lomb-Scargle periodograms were implemented using SciPy version 1.7, and cosinor analysis was implemented using CosinorPy (71). Statistical significance was defined *P < 0.05, **P < 0.01, ***P < 0.001, and n.s. for not significant.

## Acknowledgments

We thank Ryohei Narumi for the circadian liver samples and for the use of the mass spectrometry and qPCR data in (10) and Dr. Shigehiro Kuraku and GIRC at Osaka University for sequencing of the ribosome profiling libraries. We thank Dr. Shihoko Kojima and Dr. Seung-Hee Yoo for PER2::LUC MEFs. We thank all lab members at RIKEN BDR, in particular, Natsumi Hori and Yumika Sugihara for the production of mutant mice; and Masako Kunimi and Ruriko Inoue for breeding and helping phenotype measurements of the mutant mice. We thank all lab members at iFReC Systems Immunology lab and iFReC Laboratory for Host Defense, in particular, Kitiya Piboonprai for help with genotyping and Shizuo Akira for his kind support. This work was also supported by grants from Exploratory Research for Advanced Technology (ERATO) grant (JPMJER2001, H.R.U.) from the Japan Science and Technology Agency (JST), KAKENHI Grant-in-Aid from JSPS (Scientific Research S 18H05270, H.R.U.), MEXT Quantum Leap Flagship Program (MEXT QLEAP) Grant Number JPMXS0120330644, AMED under Grant Number JP20am0401011, and the intramural Grant-in-Aid from the RIKEN Center for Biosystems Dynamics Research to H.R.U. This research was supported by Research Support Project for Life Science and Drug Discovery (Basis for Supporting Innovative Drug Discovery and Life Science Research (BINDS)) from AMED under Grant Number JP22ama121025. This work was supported by the Human Frontiers Science Program and a JSPS Grant-in-Aid for Early Career Scientists (18K14755 to A.M.). Two children were born to A.M. during the course of this project, and I thank them and Mari Nishino for their care, love, and support.

## Notes

### Competing Interest Statement

The authors have declared no competing interest.

### Summary of Updates

We shifted our explanation of the Per2 uORF from a mechanism based on translation to an effect on Per2 mRNA levels and analyzed PER2 levels in nuclear and cytoplasmic lysates (Fig. S19, S20, S21). We also removed claims about period in the significance statement and provided a rationale for ribosome profiling at ZT2-4 in the text. We think these changes fully address the remaining reservations from both reviewers. Importantly, the significance of our study derives from that fact that it is the first to perform ribosome profiling under constant darkness conditions, to correlate ribosome profiling data with mass spectrometry under these conditions, and (alongside the work by Miyake et al.) to identify and characterize the uORF in Per2. We firmly believe that these novel contributions significantly advance the field of circadian research and provide compelling reasons for its publication.

