## Supplementary material for "Circadian ribosome profiling reveals a role for the *Period2* upstream open reading frame in sleep": SI_Appendix

PNAS

**This PDF file includes:**

- Supporting text
- Figures S1 to S21
- Tables S1 to S3
- Legends for Datasets S1 to S5
- SI References

**Other supporting materials for this manuscript include the following:**

- Datasets S1 to S5

**Supporting Information Text**

### Methods

**Animals.** *Per2* uORF mutant (uORF: ATGTGA to mutant uORF: GTAGGT) mice were generated by one-cell embryo microinjection of synthesized Cas9 mRNA, gRNA (5'-TTTCCACTATGTGACAGCGGAGG-3'), and ssODN (5'-GTCACGTTTTCCACTGTAGGTCAGCGGAGGGCGACG-3') in C57BL/6N fertilized eggs, and genotypes were confirmed by Sanger sequencing.

**Plasmids.** The pGL3-P(*Per2*)-d*Luc* reporter plasmid (1) was modified by standard PCR mutagenesis to generate pGL3-P(*Per2*)-uORF-d*Luc* (ATG->TTG mutation in the *Per2* uORF). Synthetic oligonucleotide linkers containing 0-4 artificial uORFs (Fastmac, Japan) were inserted into a PCR-linearized pGL3-P(*Per2*)-d*Luc* and pGL3-P(SV40)-3xE'box-d*Luc* (2) by In-Fusion cloning (Takara Bio). Plasmids containing full-length *Per2* 5'UTR and PER2 were cloned from mouse cDNA (Clone PX00938 H17, RIKEN) and pMU2-PER2-Luc (3), respectively, and inserted into pGL3-P(*Per2*)-d*Luc* or pGL3-P(*Per2*)-uORF-d*Luc* by In-Fusion cloning to create pGL3-P(*Per2*)-5'UTR-d*Luc* and pGL3-P(*Per2*)-5'UTR-PER2-d*Luc* and the respective uORF mutant plasmids. Oligos for *Per2* sgRNA (5'-caccgTTCCACTATGTGACAGCGGA and 5'-aaacTCCGCTGTCACATAGTGGAAc) were annealed and inserted into *Bbs*I-digested pSpCas9(BB)-2A-Puro (Addgene #62988) or pSpCas9n(BB)-2A-Puro (Addgene #62987) to make pCas9-Per2 or pCas9n-Per2, respectively. The promoter sequences from pGL3-P(*Per2*)-d*Luc* or pGL3-P(*Per2*)-uORF-d*Luc* were cloned into pD1EGFP-N1 (Takara Bio) to create pD1-P(*Per2*)-EGFP or pD1-P(*Per2*)-uORF-EGFP, respectively. mCherry was PCR-amplified and blunt-end ligated into inverse-PCR amplified pD1-P(*Per2*)-EGFP for the mCherry version. The *Nr1d1* 5'UTR (nucleotides 1-629 in NM_145434.4) and *Cry1* 5'UTR (nucleotides 1-583 in NM_007771.3) were inserted into phpRL-BCL2-FL-pA (Addgene #42595) by In-Fusion cloning to create phpRL-*Nr1d1* 5'UTR-pA and phpRL-*Cry1* 5'UTR-pA, respectively. All plasmid sequences were verified by Sanger sequencing.

**Ribosome Profiling.** RNA was precipitated from the elution by the addition of 38.5 µl water, 1.5 µl GlycoBlue (ThermoFisher), and 10 µl 3 M sodium acetate pH 5.5 followed by 150 µl isopropanol, stored overnight at -80 °C, pelleted by centrifugation (20,000 × *g* for 30 min at 4 °C), and resuspended in 5 µl 10 mM Tris pH 8. Approximately 2–4 µg RNA was separated on a 15% (wt/vol) polyacrylamide TBE-urea gel at 200 V for 65 min in 1× TBE, and fragments between 26 and 34 nt were excised and recovered in 400 µl RNA gel extraction buffer (300 mM sodium acetate pH 5.5, 1 mM EDTA, and 0.25% (wt/vol) SDS) overnight. RNA was precipitated by the addition of 1.5 µl GlycoBlue and 500 µl, stored at -80 °C for >30 min, pelleted by centrifugation (20,000 × *g* for 30 min at 4 °C), and resuspended in 10 µl 10 mM Tris pH 8. RNA was dephosphorylated by T4 PNK (Takara Bio), precipitated and recovered as previously described, resuspended in 8.5 µl 10 mM Tris, and 1.5 µl of preadenylated and 3' blocked miRNA cloning linker (1/5rApp/CTGTAGGCACCATCAAT/3ddC/, IDT) was added. The linker mixture was denatured for 90 s at 80 °C, cooled to room temperature, ligated by T4 Rnl2 (NEB) for ~5 h at room temperature, precipitated, recovered, and separated by 15% TBE-urea gel electrophoresis. Ligation products were recovered overnight in RNA gel extraction buffer, precipitated, and recovered in 10 µl 10 mM Tris pH 8. Ligated RNA was reverse transcribed at 48 °C for 30 min using SuperScript III (ThermoFisher) with the reverse transcription primer (5′-(Phos)-AGATCGGAAGAGCGTCGTGTAGGGAAAGAGTGTAGATCTCGGTGGTCGC-

(SpC18)-CACTCA-(SpC18)- TTCAGACGTGTGCTCTTCCGATCTATTGATGGTGCCTACAG-3′, where SpC18 indicates a hexa-ethyleneglycol spacer). Reverse transcription products were precipitated, recovered, separated by 15% TBE-urea gel electrophoresis, and recovered overnight in DNA gel extraction buffer (300 mM NaCl, 10 mM Tris pH 8, and 1 mM EDTA). The cDNA was precipitated, recovered, resuspended in 15 µl 10 mM Tris pH 8, and circularized by CircLigase (EpiCentre) for 1 h at 60 °C. rRNA was depleted by combining 5 µl circularization reaction with 1 µl of subtraction oligo pool, 1 µl 20× SSC, and 3 µl water, incubating at 37 °C for 15 min and binding to MyOne Streptavidin C1 DynaBeads (ThermoFisher) at 37 °C for 15 min with mixing at 1000 rpm. Eluate was precipitated, recovered, and resuspended in 5 µl 10 mM Tris pH 8. rRNA-depleted cDNA was PCR amplified by Phusion High-Fidelity DNA Polymerase (NEB) in a 20 µl reaction (denature 94 °C for 15 s, anneal 55 °C for 10 s, extend 72 °C for 10 s) for 12-16 cycles. PCR products were separated on an 8% TBE gel, recovered overnight in DNA gel extraction buffer, precipitated, recovered, and resuspended in 15 µl 10 mM Tris pH 8. Libraries were quantified using the high-sensitivity DNA chip on the Agilent BioAnalyzer according to the manufacturer's protocol, pooled, and amplified on the Illumina HiSeq or NovaSeq6000 system according to the manufacturer's protocol.

**Bioinformatic analysis of ribosome profiling.** After quality control using FASTQC (https://www.bioinformatics.babraham.ac.uk/projects/fastqc/), reads were clipped using fastx_clipper (with parameters `-Q33 -a CTGTAGGCACCATCAAT -l 25 -c -n`) and processed with fastx_trimmer (with parameters `-Q33 -f 2`), both from the FASTX-Toolkit (http://hannonlab.cshl.edu/fastx_toolkit/). We then used Bowtie 2.1.0.0 (4) to map the resulting reads to rRNA sequences. Reads that were successfully aligned to these sequences are discarded, and only unaligned reads were used for downstream analysis. These reads that did not align to rRNA were mapped to the mouse mm10 reference genome. Using samtools 0.1.19.0 and egrep, we extracted reads for which an exact match between the sequence and the reference could be found.

Using reads aligned to a unique location, we then calculated transcript density profiles. This relies on assignment each footprint alignment (i.e., each read) to a specific A site nucleotide based on the length of the fragment. The initial assumption is that this site will be close to the center of the read. To calculate the best offset, we considered a metagene that captured all reads and their position relative to the start site and calculated the offset that optimized phasing. The best results were obtained when the offset from the 5' end of the alignment was: 26 nt long, +13; 27-28 nt long, +15; 29-30 nt long, +16; 31-32 nt long, +17. Reads shorter than 26 or longer than 32 nucleotides were discarded. Using this offset, we assigned each read to a unique nucleotide, and constructed transcript density profiles by counting the number of reads whose A site was assigned to each nucleotide position. Raw data has been deposited in GEO (GSE201732 and GSE231820).

**Cells.** PER2::LUC MEFs were plated in 10 cm dishes at a density $4 \times 10^5$ per well, transfected the next day with pCas9-Per2 or pCas9n-Per2 and 560 bp of single-stranded linearized DNA (Guide-it Long ssDNA Production System, Takara Bio) containing mutations in the Per2 uORF (ATGTGA to GTAGGT), and after 48-h selected in 1 µg/ml puromycin for puromycin-resistant clones according to (5). Individual clones were expanded, sequence-verified, and clones containing at least one allele with a mutation in the Per2 uORF were used for downstream analysis.

**Preparation of wild-type and *Per2* uORF mutant mouse lysate for ribosome profiling, total RNA sequencing, qPCR, and immunoblot analysis.** Approximately 0.1 g mouse liver tissue from wild-type and *Per2* uORF mutant male mice sacrificed at ZT02-04 was washed repeatedly in ice-cold PBS (Nacalai), resuspended in 1 ml polysome lysis buffer (150 mM NaCl, 20 mM Tris-HCl pH 7.4, 5 mM $MgCl_2$, 5 mM DTT, 100 µg/ml cycloheximide, 1% Triton X-100, 25 U/ml Turbo DNAse I), and homogenized by glass Dounce microhomogenizer (>10 strokes tip A, then >10 strokes tip B). The homogenate was clarified by centrifugation (20,000 × *g* for 10 min at 4 °C) and the supernatant divided – 300 µl was immediately used for ribosome profiling library preparation as above, 200 µl was added to 1 mL TRIzol (Nacalai) for total RNA sequencing, and the remaining was snap frozen and stored at -80 °C for subsequent western blot and qPCR analysis. Total RNA was extracted and analyzed using the Agilent 2100 bioanalyzer according to the manufacturer's instructions. Total RNA libraries were prepared using the TruSeq stranded mRNA kit (Illumina) according to the manufacturer's instructions.

**Preparation of nuclear and cytoplasmic extracts.** Extracts were prepared essentially as described previously (6) with some modifications. Approximately 0.1 g mouse liver tissue from wild-type and *Per2* uORF mutant male mice sacrificed at ZT02-04 was washed repeatedly in ice-cold PBS (Nacalai), resuspended in 1 ml per 0.1 g tissue in homogenization buffer (5 mM KCl, 10 mM Tris-HCl pH 7.4, 2 mM EDTA, 1 mM DTT, 1× cOmplete Mini EDTA-free protease inhibitor tablet [Sigma]) + 2.2 M sucrose, and homogenized by glass Dounce microhomogenizer (>10 strokes tip A, then >10 strokes tip B). Approximately 550 µl lysate was layered on 1.5 ml sucrose cushion (homogenization buffer + 2.2 M sucrose + 10% glycerol) in 13 mm × 51 mm polycarbonate ultracentrifuge tubes, and centrifuged in a TLA100.3 rotor at 70,000 rpm at 4 °C for 45 min. The supernatant was removed and used as the cytoplasmic extract. The nuclear pellet was rinsed once glycerol without disturbing the pellet with homogenization buffer + 10% glycerol, then resuspended in 150 µl homogenization buffer + 10% glycerol, transferred to a new tube containing 150 µl urea buffer (2 M urea, 600 mM NaCl, 50 mM Tris pH 7.4, 1 mM DTT, 1× cOmplete Mini EDTA-free protease inhibitor tablet [Sigma]), incubated on ice for 20 min with occasional mixing, and then centrifuged (20,000 × *g* for 10 min at 4 °C). This supernatant was used as the nuclear extract.

**Western blot.** Total protein lysates were thawed and diluted to ~10 mg/ml in polysome lysis buffer. Cytoplasmic and nuclear lysates were diluted to 1 mg/ml and 0.6 mg/ml, respectively, in homogenization buffer + 10% glycerol. Lysates were separated by SDS-PAGE, transferred to nitrocellulose membrane (Thermofisher) by semidry transfer (Bio-Rad), blocked in 5% (wt/vol) skim milk in TBST (50 mM Tris pH 7.5, 140 mM NaCl, 0.1% Tween) for 1 h at 22 °C, and then incubated overnight at 4 °C in anti-PER2 antibody (1:1000, PM083, Medical & Biological Laboratories) or anti-histone H3 antibody (1:1000, 9715, Cell Signaling). After washing in TBST, blots were incubated in anti-rabbit IgG HRP-conjugated secondary antibody (1:2000, NA934, Cytivia) or mouse beta-actin conjugated-HRP antibody (0.8:2000, sc-47778, Santa Cruz Biotechnology) for 1 h at 22 °C and developed by Immobilon Forte Western HRP (Millipore).

**qPCR.** Approximately 100 µl lysate was extracted in 1 ml TRIzol (Nacalai) according to the manufacturer's instructions and re-suspended in 50 µl 10 mM Tris pH 8.0. RNA was digested with TURBO DNAseI (Thermofisher) for 1 h at 37 °C and recovered with an equal volume 1:1 phenol:chloroform containing isoamyl alcohol pH 5.2 (Nacalai). After centrifugation (20,000 × *g* for 5 min at 4 °C), approximately 37.5 µl of the aqueous layer was precipitated with 10 µl 3 M sodium acetate pH 5.5 and 1.5 µl GlycoBlue followed by 150 µl isopropanol. Samples were frozen at -80 °C for >30 min, centrifuged (20,000 × *g* for 30 min at 4 °C), washed in 500 µl 75% ethanol, dried for 10 min at 37 °C, and RNA was resuspended in 15 µl 10 mM Tris pH 8.0. Approximately 500 ng RNA was reverse transcribed using ReverTra Ace qPCR RT Master Mix (Toyobo) and amplified using Thunderbird SYBR qPCR mix (Toyobo) with *Per2* (forward: 5'-GCACATCTGGCACATCTCGG-3'; reverse: 5'-TGGCATCACTGTTCTGAGTGTC-3') and *actin* (forward: 5'-CACTGTCGAGTCGCGTCCA-3'; reverse: 5'-CATCCATGGCGAACTGGTGG-3') primers on ViiA 7 (Applied Biosystems). Relative *Per2* mRNA was quantified using the ΔΔCT method with *actin* as a reference gene.

**Sleep and wake parameters.** Sleep and wake duration are the mean total sleep and wake duration, respectively, per day over 13 d. $P_{sw}$ is the transition probability from sleep to awake; $P_{ws}$ is the transition probability from awake to sleep. Mathematically, $E_i$ is the *i*-th epoch, which is either sleep (*s*) or wake (*w*). Let $N_{XY}$ ($X,Y \in \{w,s\}$) be the number of elements of the set $\{(E_i, E_{i+1}) \mid E_i = X, E_{i+1} = Y\}$. $P_{ws}$ and $P_{sw}$ are defined as $N_{ws}/(N_{ws} + N_{ww})$ and $N_{sw}/(N_{sw} + N_{ss})$, respectively. The amplitude is defined as the coefficient of variation of sleep time for each 10-min bin over 24 h.

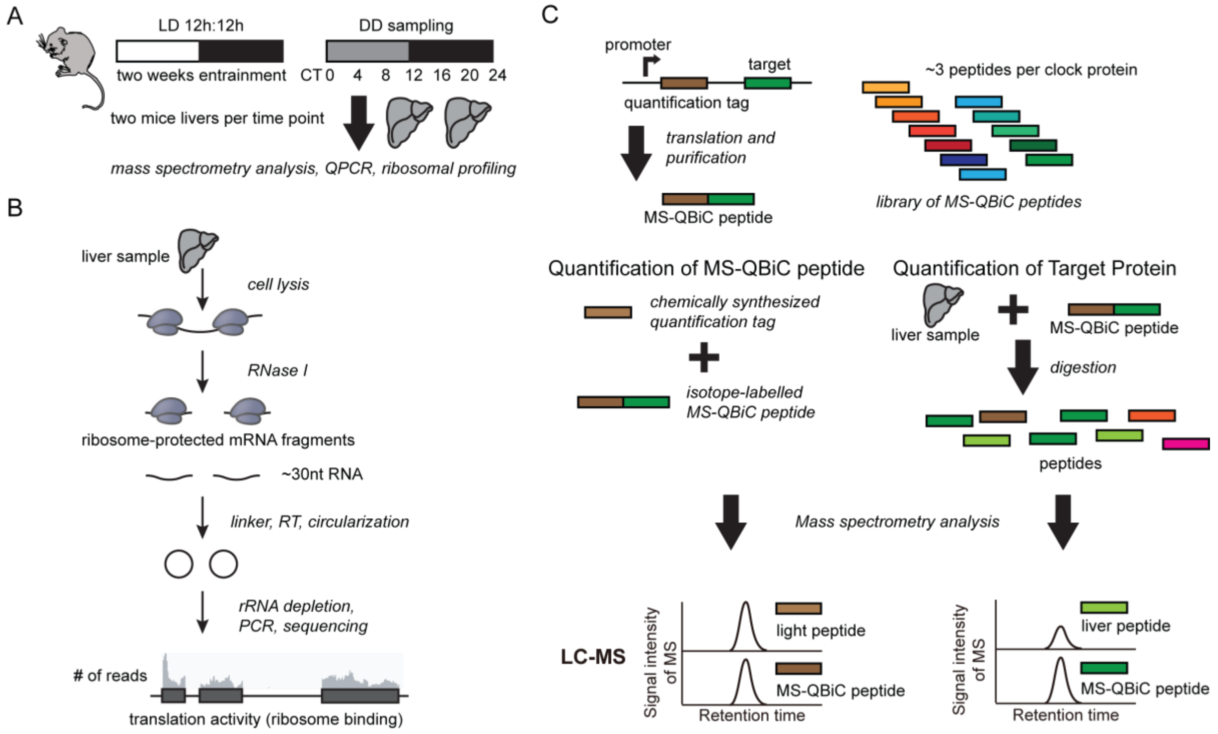


**Supplementary Figure 1 | Analyzing mRNA expression, translation activity, and absolute protein copy number levels in the same mice liver samples over the course of a day (related to Figure 1). (A)** All mice were carefully kept and handled according to the RIKEN Regulations for Animal Experiments. Eight to ten-week-old-male mice (C57BL/6N, Japan SLC) were entrained under a 12–12 h light-dark (LD) conditions (400 lux) for 2 weeks. After transferring to constant darkness (DD), mice were sacrificed every 4 h over 1 day (i.e. CT0, 4, 8, 12, 16, and 20) for proteomic analyses, every 4 h over 1 day (i.e. CT0, 4, 8, 12, 16, 20, and 24) for ribosome profiling analyses, or every 4 h over 2 days (i.e. CT0, 4, 8, 12, 16, 20, 24, 28, 32, 36, 40, and 44) for qPCR analyses. **(B)** Two mice livers at seven different circadian time points were lysed and ribosomes were isolated by a sucrose cushion. Ribosome-protected mRNA fragments were extracted, reverse transcribed, and circularized into a library. Libraries were depleted of the common most abundant rRNA, PCR amplified and sequenced (~1.02 billion reads). **(C)** Schematic description of the MS-QBiC workflow. A purification-tag, a quantification-tag, and a tryptic peptide of the target protein (target peptide) were sequentially arrayed as a single peptide sequence (MS-QBiC peptide). The target peptide sequence was attached by one- or two-step PCR. The MS-QBiC peptide was synthesized in the PURE system in the presence of stable isotope-labeled Arg and Lys for isotopic labeling both the quantification-tag and the target peptide. Trypsin digestion of purified MS-QBiC peptide produced equal amounts of isotopically labeled quantification-tag and target peptide. The quantification-tag was used to measure purified MS-QBiC peptide and the target peptide was used as an internal standard for target protein quantification.

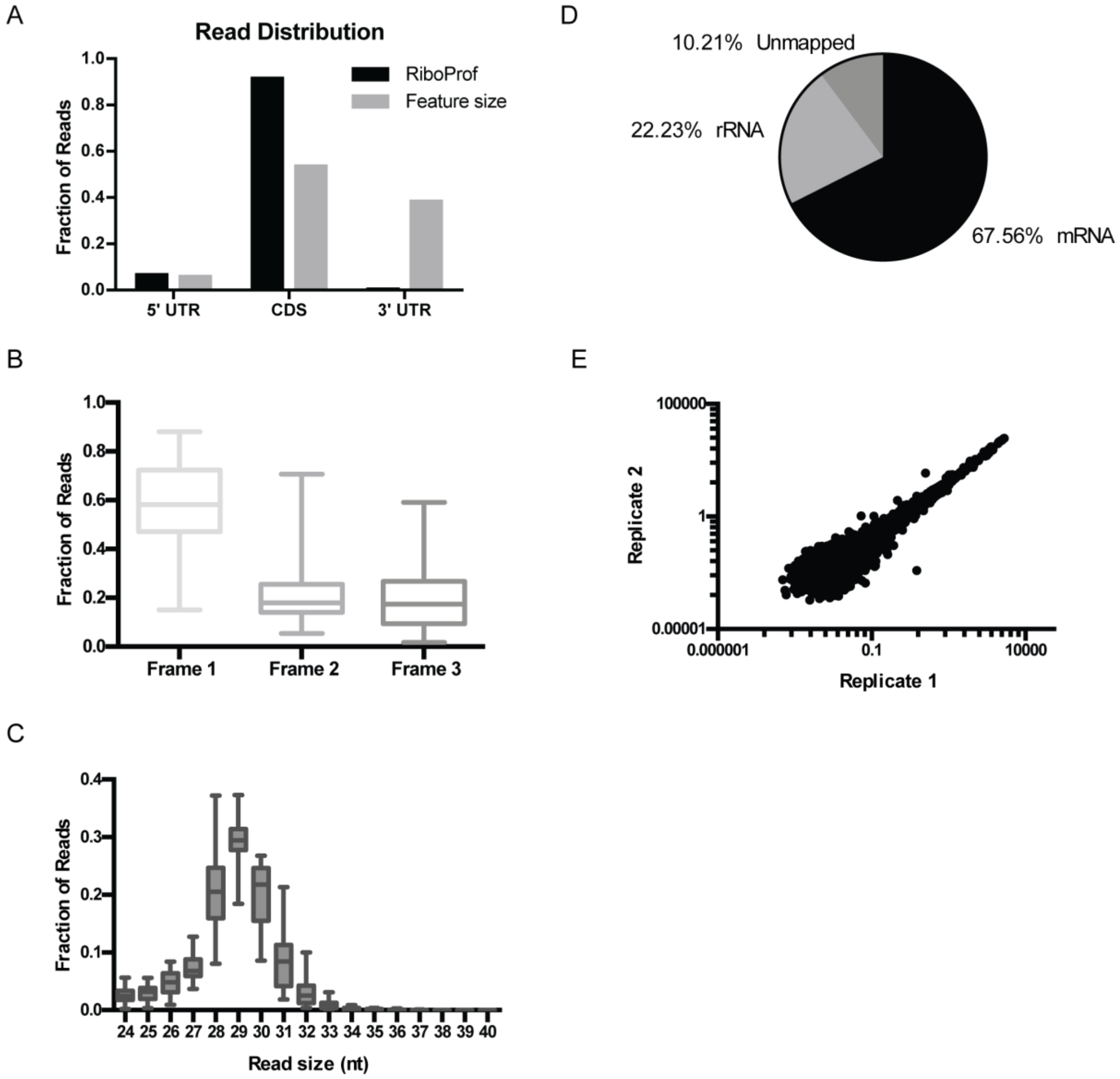


**Supplementary Figure 2 | Validation of ribosome profiling cDNA libraries (related to Figure 1). (A)** Read distribution within 5'UTRs, CDS, and 3'UTRs compared with the size of those features. Ribosome profiling reads are enriched for the CDS and 5'UTR, and relatively few reads map to the 3'UTR. **(B)** Frame analysis of ribosome profiling reads of single protein isoform mRNAs shows a preference for frame 1 (the coding frame). Box-and-whisker plots: midline, median; box, 25th and 75th percentiles. Whiskers are minimum and maximum values. **(C)** Box and whisker plots show the distribution of read lengths from all sequences. The majority of reads were between 27 and 32 nt, which matches the footprint size of the ribosome. **(D)** Summary of mapped reads. **(E)** Correlation of RPKM between replicate 1 and replicate 2 at CT0 shows a high degree of correlation in ribosome binding between samples.

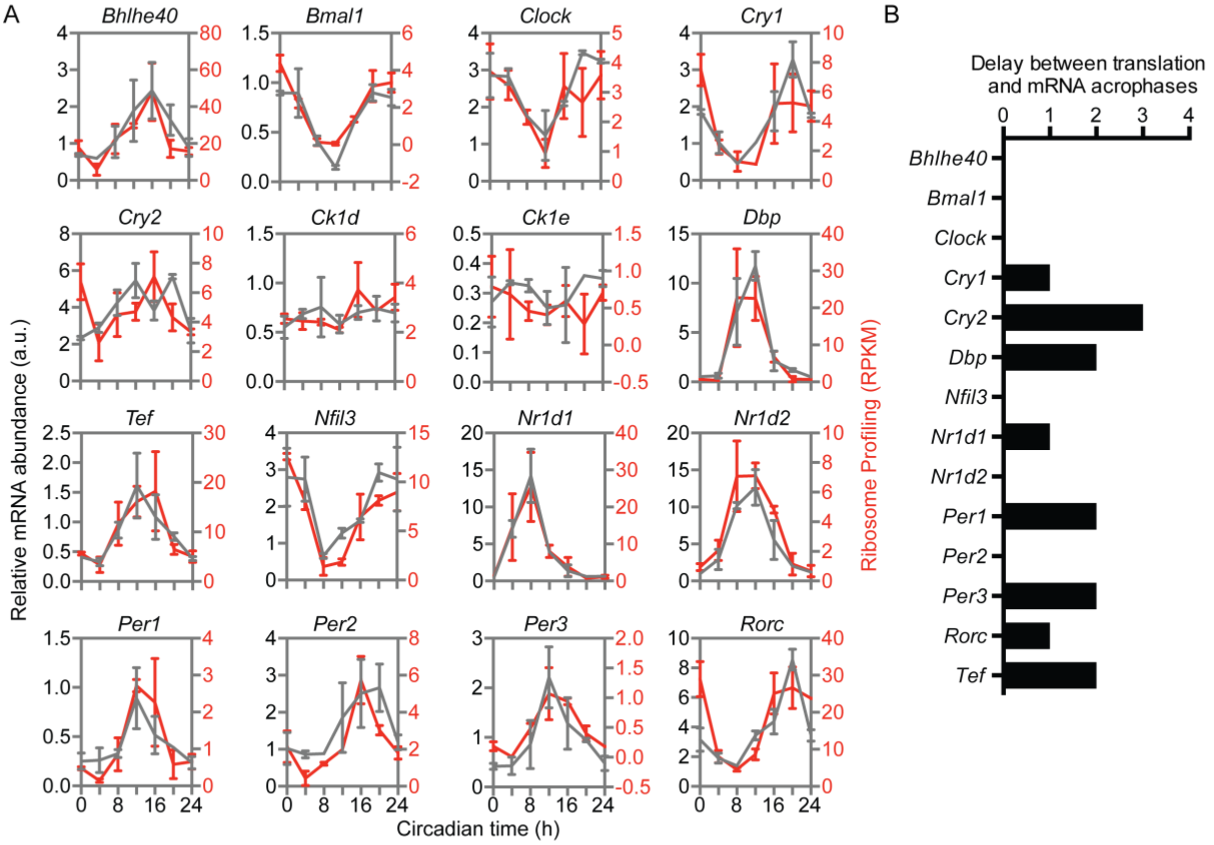


**Supplementary Figure 3 | Measuring translation and RNA expression from the same liver samples for 16 circadian mRNAs (related to Figure 1). (A)** Time course of the quantified RNA expression levels (grey) by qPCR derived from data in (7) and the corresponding ribosome binding (RPKM, red). Error bars are the standard deviation of the two replicates. **(B)** Phase delay between mRNA expression and ribosome binding. Absolute cDNA abundance was calculated using the standard curve obtained from mouse genomic DNAs, and JTK analysis was used to estimate the phase of each mRNA.

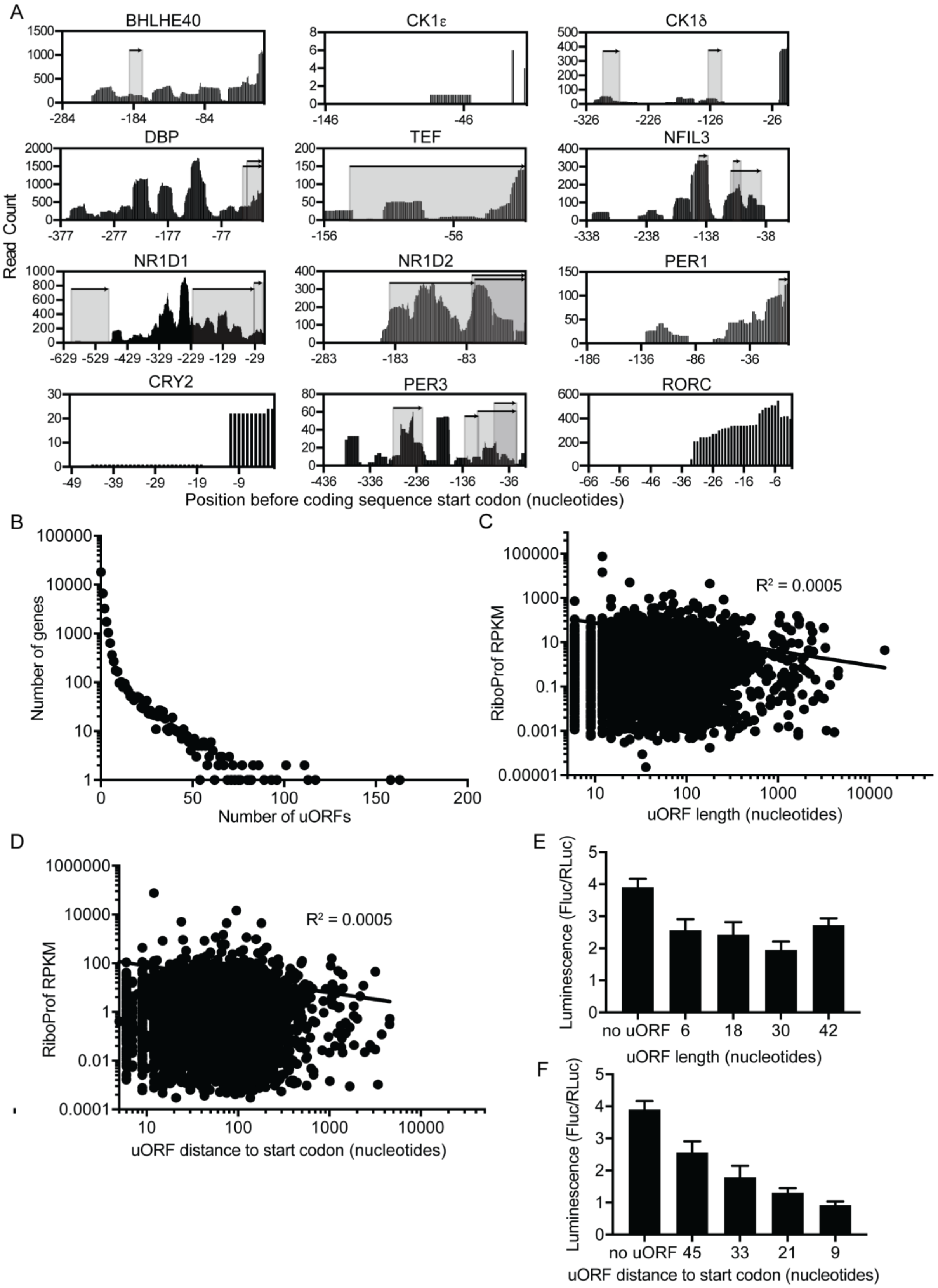


**Supplementary Figure 4 | Distribution of upstream open reading frames in clock transcripts (related to Figure 2). (A)** The 5' UTRs of *Bhlhe40, Ck1e, Ck1d, Dbp, Tef, Nfil3, Nr1d1*, *Nr1d2*, *Per1*, *Cry2*, *Per3,* and *Rorc* transcripts, location of uORFs (shaded regions), and raw read counts from the ribosome profiling data (black bars). **(B)** The number of transcripts with a given number of uORFs. **(C)** The relationship between the length of an uORF and ribosome binding (RPKM) in the corresponding ORF. **(D)** The relationship between the distance of an uORF to the ORF start codon and ribosome binding (RPKM) in the corresponding ORF. **(E)** Introduction of a single uORF in the *Per2* short promoter of different lengths does not alter the relative luminescence of the *Per2* luciferase reporter. **(F)** The distance of a single uORF in the *Per2* short promoter to the start codon correlates with the relative luminescence of the *Per2* luciferase reporter.

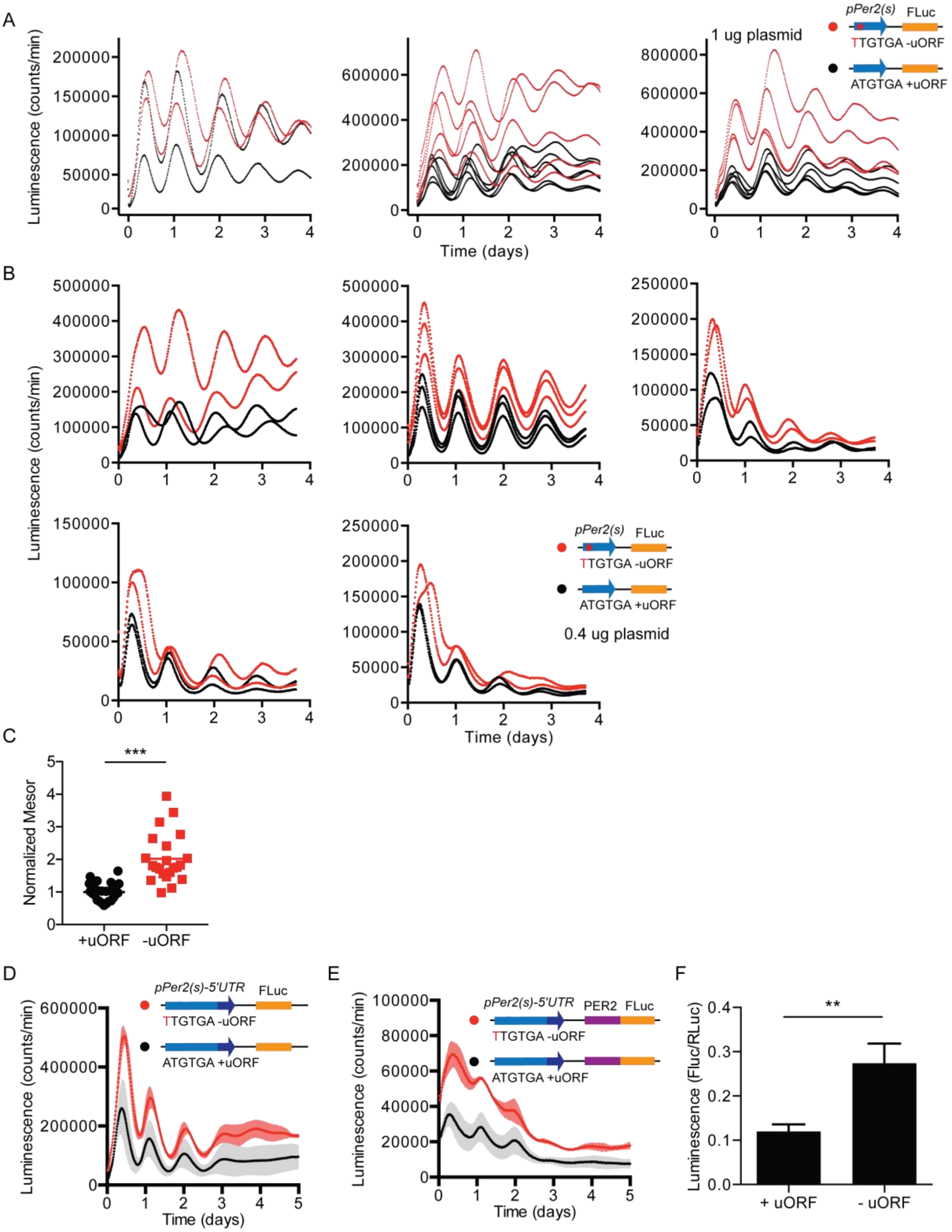


**Supplementary Figure 5 | Detailed analysis of the *Per2* uORF (related to Figure 3). (A)** Bioluminescent recording of 3T3 cells transfected with the *Per2* short promoter containing a wild-type (black) or mutant (red) *Per2* uORF at 1 µg (**A**) and 0.4 µg (**B**) transfected plasmid. Each graph is an independent experiment performed on a different day; multiple traces on the same graph indicate different transfected plates. Data from the upper middle panel of (**B**) was used for Fig. 3B. **(C)** Cosinor analysis showing normalized mesor for the bioluminescent traces in (**A**) and (**B**). **(D)** Mutation of the *Per2* uORF (ATGTAA to TTGTGA) with the full-length *Per2* 5' UTR (red) increases the luminescence expression level of the reporter as measured by bioluminescent recording compared to that of a reporter containing the uORF (black). Shaded region is SD. **(E)** Mutation of the *Per2* uORF (ATGTAA to TTGTGA) with the full-length *Per2* 5' UTR and PER2 protein (red) increases the luminescence expression level of the reporter as measured by bioluminescent recording compared to that of a reporter containing the uORF (black). Shaded region is SD. **(F)** Relative luminescence experiments of 3T3 cells transfected with (+uORF) or without (-uORF) the uORF in pGL3-P(*Per2*)-d*Luc*.

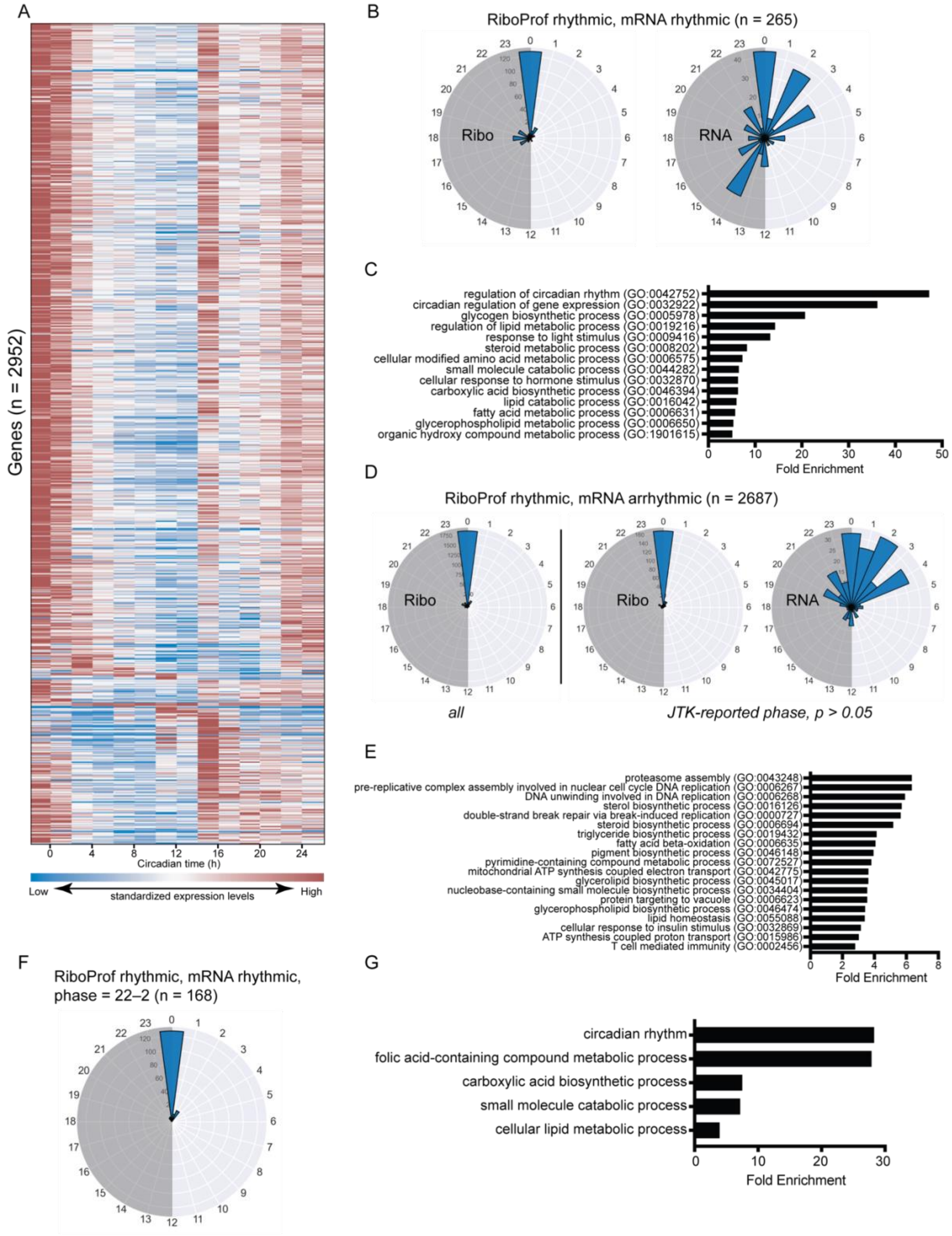


**Supplementary Figure 6 | Phase analysis (related to Figure 3). (A)** Heat map of ribosome profiling rhythmic transcripts identified by JTK cycling analysis (P < 0.05) showing RPKM data over 24 hours for 2952 genes. Transcripts are sorted by phase, and RPKM values are normalized within each gene (row). Note that most transcripts peak at CT0 (red bars in first two columns). **(B)** Polar histogram showing the ribosome profiling (*left*) and RNA (*right*)

phase distribution of the 265 ribosome profiling rhythmic and mRNA rhythmic transcripts. **(C)** Panther analysis of gene ontology biological processes based on genes in (**B**) showing an increase in circadian rhythm regulation pathways and metabolic processes. **(D)** Polar histogram showing the ribosome profiling phase distribution (*left*) of the 2687 ribosome profiling rhythmic transcripts that did not reach significance in mRNA rhythmicity. Ribosome profiling (*middle*) and RNA (*right*) polar histograms for transcripts with an mRNA JTK-reported phase $P > 0.05$ are also shown. **(E)** Panther analysis of gene ontology biological processes based on the 2687 genes in (**D**). (**F** and **G)** There were 168 transcripts from (**B**) with a phase distribution between CT22-CT0 (**F**) corresponding to circadian rhythm and metabolism processes by Panther analysis (**G**). Note that this group includes NR1D1-regulated transcripts including *Npas2*, *Arntl*, *Cry1*, *Nfil3*, and *Clock*.

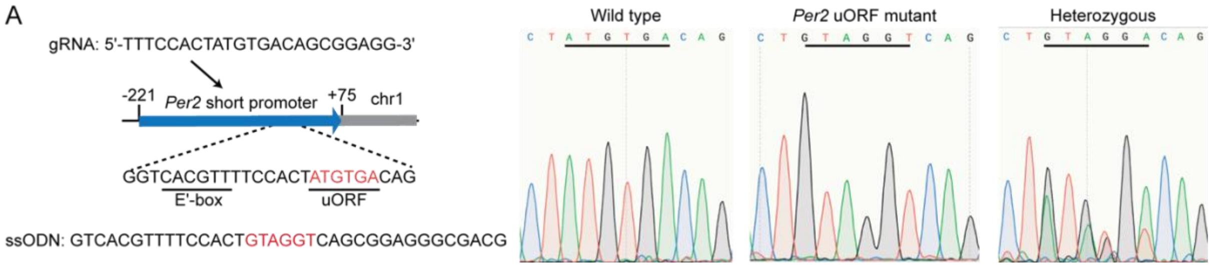


**Supplementary Figure 7 | Disruption of the *Per2* uORF in mice (related to Figure 4). (A)** An ssODN replaced the *Per2* uORF (ATGTGA) with a mutant version (GTAGGT) in which both the start and stop codons were mutated. The diagram shows the location of the CRISPR gRNA and upstream E'box within the *Per2* promoter region (*left*). Representative Sanger sequencing results from wild-type, *Per2* uORF mutant, and heterozygous mice are shown (*right*).

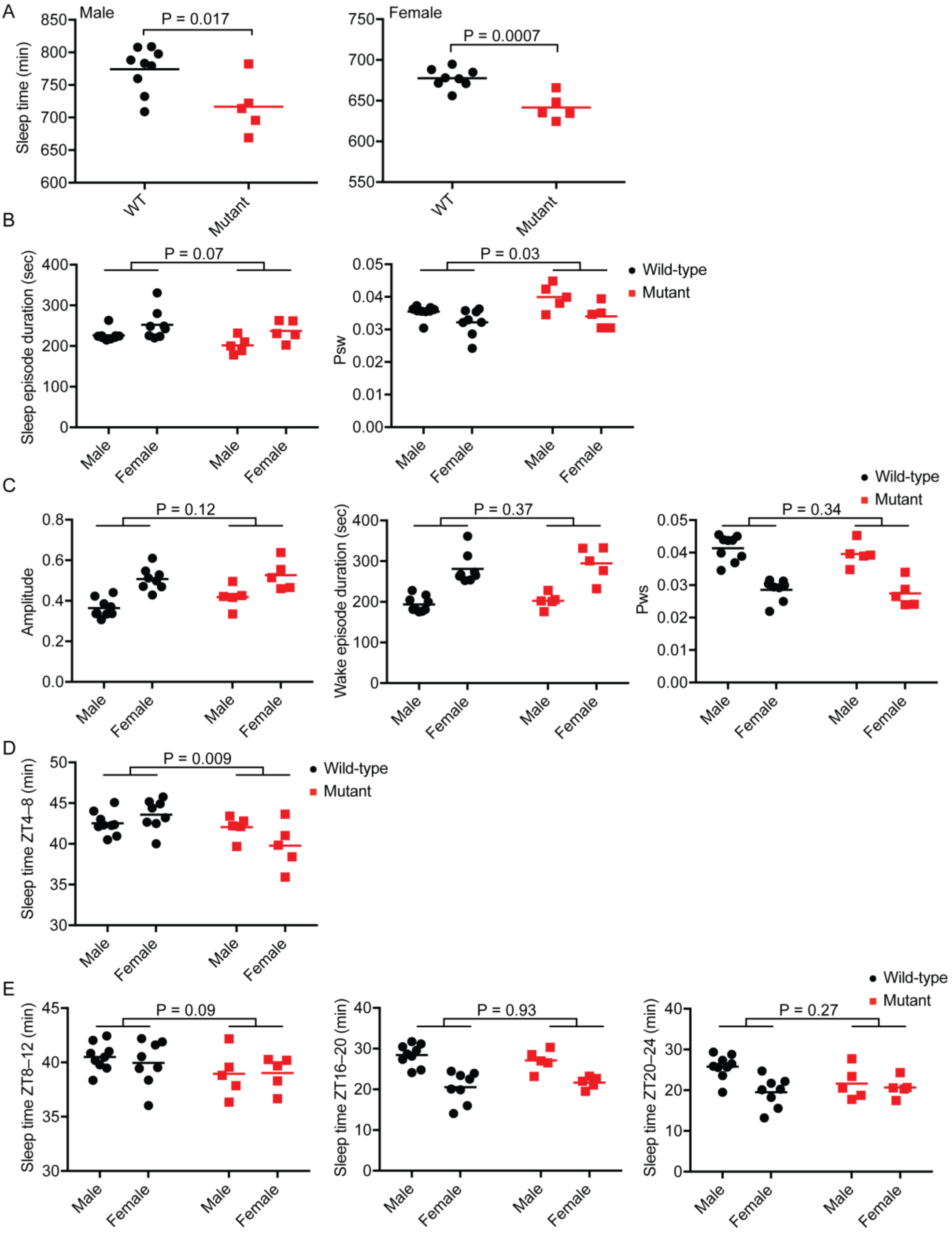


**Supplementary Figure 8 | Detailed analysis of 12 h light:12 h dark (LD) sleep parameters (related to Figure 4). (A)** Mean sleep duration over 24 h averaged over 13 d for male (*left*) and female (*right*) mice. Red, *Per2* uORF mutant mice. Black, wild-type. P values, Student's unpaired t-test. **(B)** Sleep episode duration (*left*) was shorter and $P_{sw}$ was higher (*right*) in *Per2* uORF mutant mice than in wild-type mice, but the differences were not significant. **(C)** There were no significant differences in amplitude (*left*), wake episode duration (*middle*), and $P_{ws}$ (*right*) between *Per2* uORF mutant and wild-type mice. **(D** and **E)** Mean sleep duration per hour averaged over 13 d for 4-h windows from ZT4–8 (**D**) and ZT8–12, ZT16–20, and ZT20–24 (**E**).

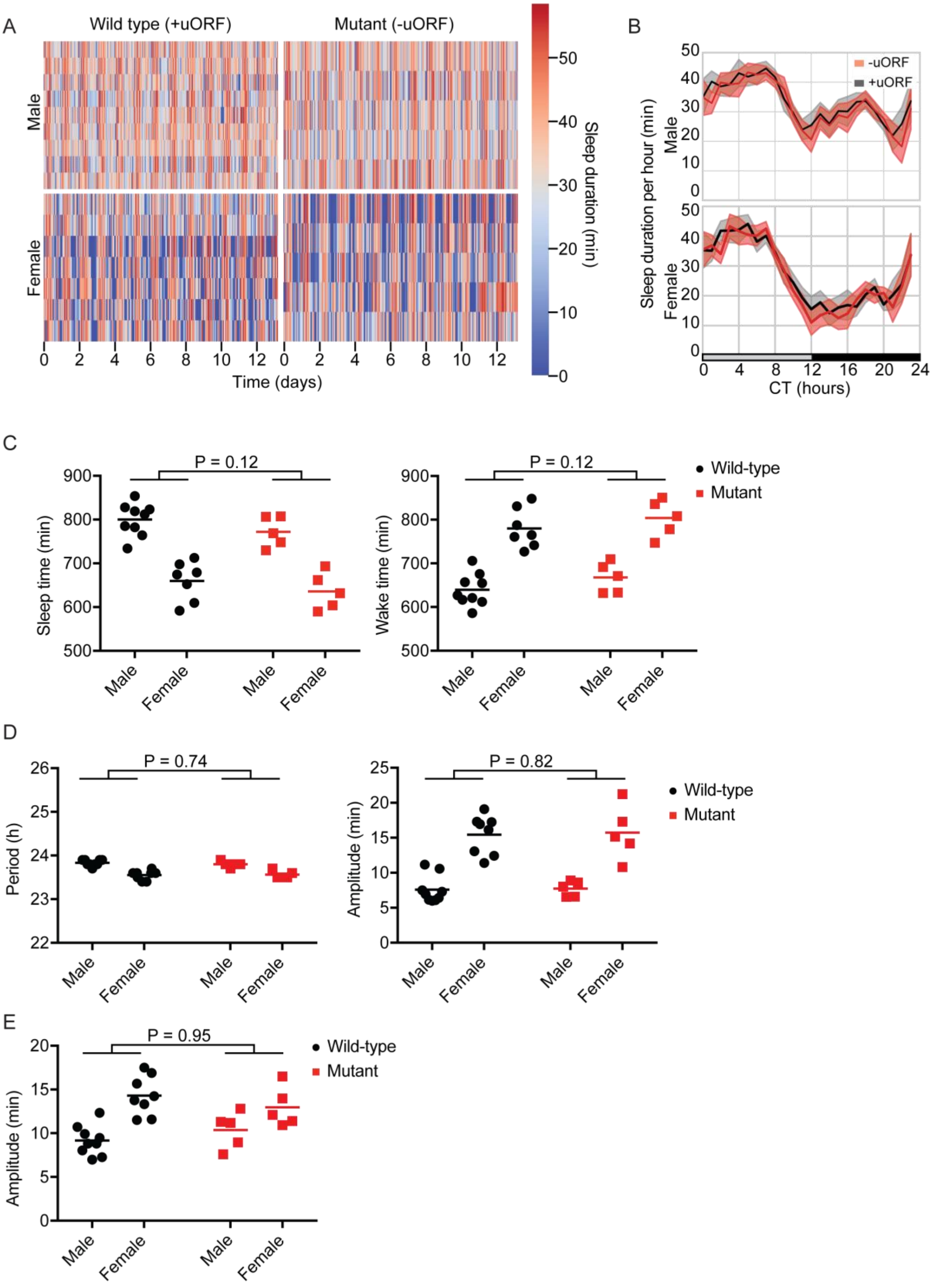


**Supplementary Figure 9 | Analysis of mice in DD conditions, and cosinor analysis of LD and DD circadian rhythm parameters (related to Figure 4). (A)** Sleep duration per hour over 12 d in constant darkness (DD) conditions for *Per2* uORF mutant and wild-type male and female mice. Each row indicates data from one mice. **(B)** Sleep duration per hour over 24 h in DD, averaged over 12 d for *Per2* uORF mutant (red) and wild-type (black) male (*top*) and female (*bottom*) mice. Lines indicate mean sleep duration at each time of day for each strain. Shaded area, SD at each time point. **(C)** Mean sleep (*left*) and wake (*right*) duration over 24 h, averaged over 12 d. Red, *Per2* uORF mutant mice. Black, wild-type. **(D)** Cosinor analysis of the period (*left*) and amplitude (*right*) of wild-type and *Per2* uORF mutant mice in DD conditions. **(E)** Cosinor analysis of the amplitude of wild-type and *Per2* uORF mutant mice in LD conditions.

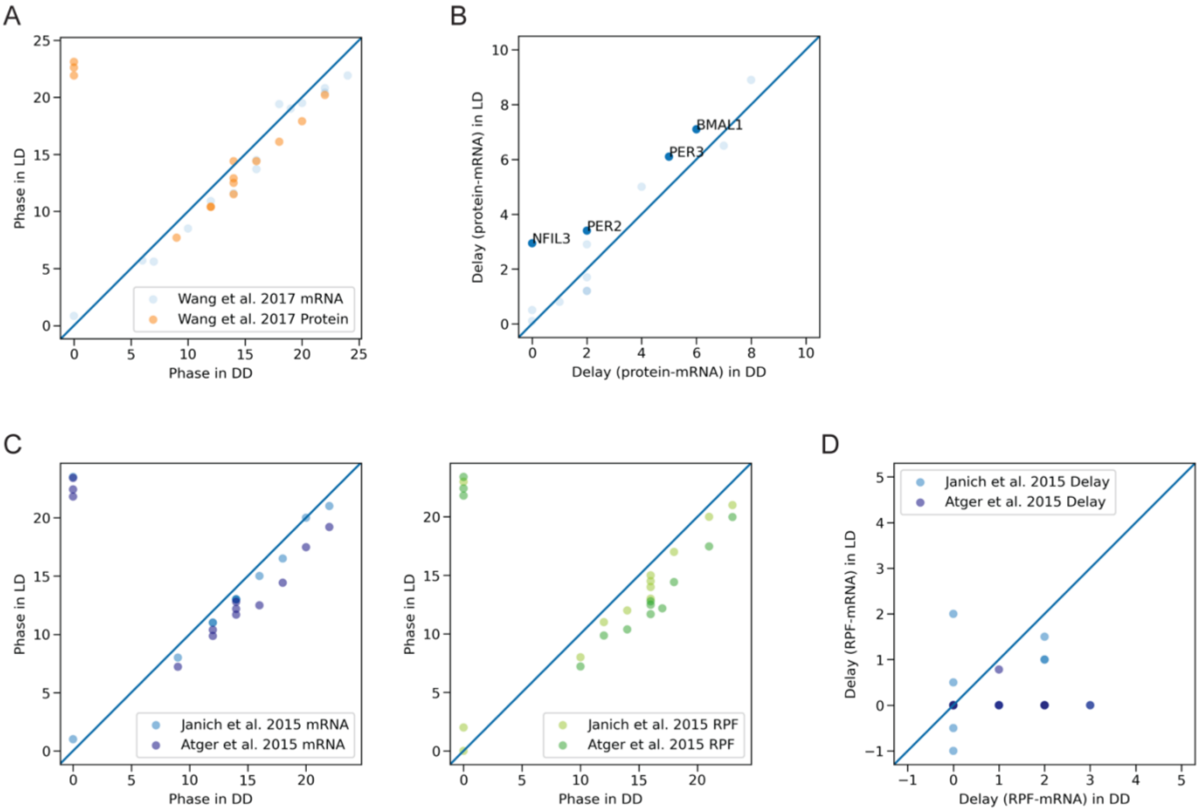


**Supplementary Figure 10 | Comparison of RNA expression, translation, and protein production phases in DD (this study) to reported phases from previously published studies in LD conditions (6, 8, 9). (A)** Protein (orange) and mRNA (blue) phases for 16 selected circadian genes observed in DD (this study) or LD (6). **(B)** The delay between mRNA and protein phases observed in DD (this study) or LD (6). Several proteins (labeled) including PER2 had a greater than one hour difference in protein-mRNA delay between LD and DD conditions. **(C)** The mRNA (*left*) and ribosome profiling (*right*) phases for 16 selected circadian genes observed in DD (this study) or LD (8, 9). **(D)** The delay between mRNA and ribosome profiling phases observed in DD (this study) or LD (8, 9).

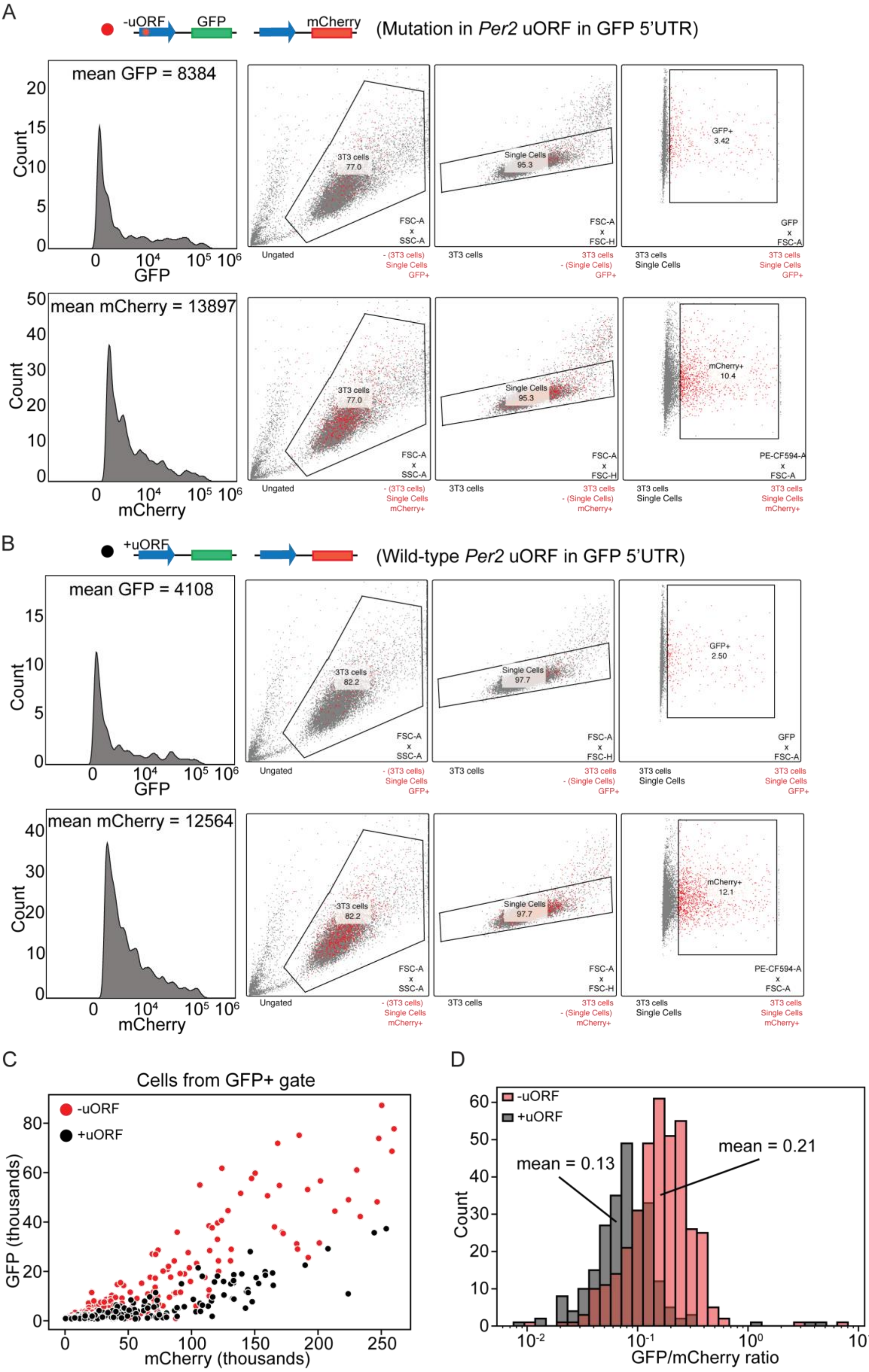
A
-uORF
GFP
mCherry
(Mutation in Per2 uORF in GFP 5'UTR)
mean GFP = 8384
Count
GFP
3T3 cells
77.0
Single Cells
95.3
GFP+
3.42
FSC-A x SSC-A
FSC-A x FSC-H
GFP x FSC-A
Ungated
mean mCherry = 13897
mCherry
mCherry+
10.4
PE-CF594-A x FSC-A
B
+uORF
(Wild-type Per2 uORF in GFP 5'UTR)
mean GFP = 4108
82.2
97.7
GFP+
2.50
mean mCherry = 12564
mCherry+
12.1
C
Cells from GFP+ gate
-uORF
+uORF
GFP (thousands)
mCherry (thousands)
D
mean = 0.13
mean = 0.21
GFP/mCherry ratio

**Supplementary Figure 11 | Extended FACS analysis of fluorescent reporter cells with and without the *Per2* uORF (related to Figure 3). (A)** Histogram (*left*) of GFP+ (top) and mCherry+ (*bottom*) cells from 3T3 cells transfected with pD1-P(*Per2*)-uORF-EGFP (-uORF, red) and pD1-P(*Per2*)-mCherry. **(B)** Histogram (*left*) of GFP+ (*top*) and mCherry+ (*bottom*) cells from 3T3 cells transfected with pD1-P(*Per2*)-EGFP (+uORF, black) and pD1-P(*Per2*)-mCherry. The backgating strategies for each gate in (**A)** and (**B**) are shown (*right*), including the percentage of GFP+ and mCherry+ cells in the population (*rightmost graph*). **(C)** Scatterplot of GFP versus mCherry expression for -uORF (red) and +uORF (black) cells gated on GFP+ expression. **(D)** Histogram of the GFP/mCherry ratio for cells in (**C**).

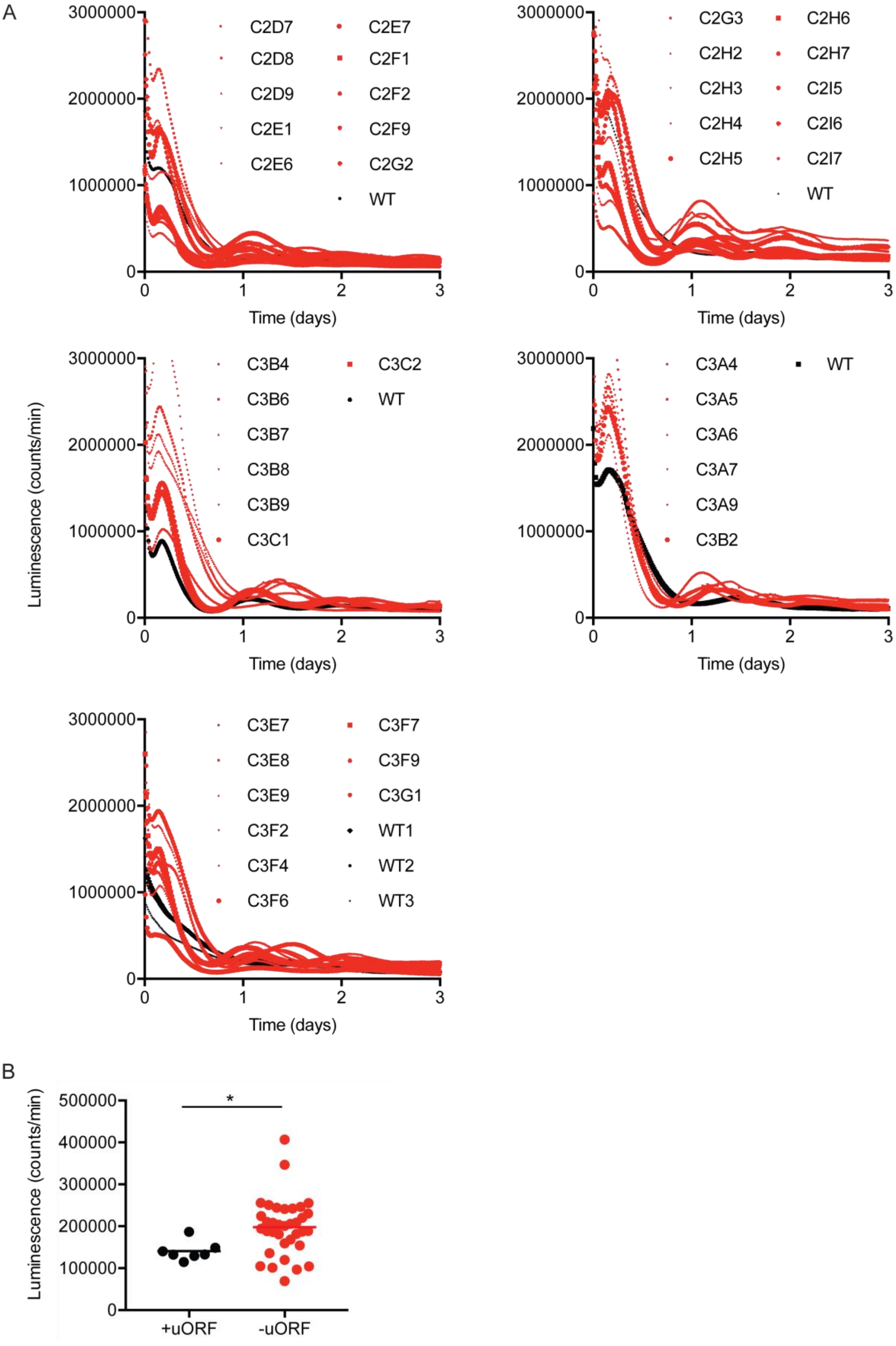
A
3000000
2000000
1000000
0
0
1
2
3
Time (days)
C2D7
C2D8
C2D9
C2E1
C2E6
C2E7
C2F1
C2F2
C2F9
C2G2
WT
C2G3
C2H2
C2H3
C2H4
C2H5
C2H6
C2H7
C2I5
C2I6
C2I7
WT
Luminescence (counts/min)
C3B4
C3B6
C3B7
C3B8
C3B9
C3C1
C3C2
WT
C3A4
C3A5
C3A6
C3A7
C3A9
C3B2
WT
C3E7
C3E8
C3E9
C3F2
C3F4
C3F6
C3F7
C3F9
C3G1
WT1
WT2
WT3
B
Luminescence (counts/min)
500000
400000
300000
200000
100000
0
*
+uORF
-uORF

1

**Supplementary Figure 12 | Detailed analysis of the *Per2* uORF in PER2:LUC MEF cells. (A)** Bioluminescent recordings of wild-type PER2:LUC MEF cells (black) or heterozygous PER2:LUC MEF clones containing a mutated *Per2* uORF (ATGTGA to GTAGGT,  red). Each graph is an independent experiment performed on a different day; multiple traces on the same graph indicate different clones. **(B)** Average bioluminescence expression between 24 h and 72 h after dexamethasone stimulation of wild-type (black) and *Per2* uORF mutant (red) MEFs.

A

```
CLUSTAL format alignment by MAFFT (v7.505)

                                               E' Box      Per2 uORF
Mouse           ---------------------gcgcgcggtcacgttttccactatgtgacagcggagggc
Rat             gcgcggaggggcgggctcaacgcgcgcggtcacgttttccactatgtgacagcggagggc
GuineaPig       ---------------------gcgcgcggtcacgttttccactatgtgacagcggcgacc
Human           ----------------------------------------actatgtgacagcggcgact
Rhesus          ------------------ggcgcgcgcggtcacgttttccactatgtgacagcggcgact
Marmoset        ------------gggcgcagcgcgcgcggtcacgttttccactatgtgacagcggcgact
Rabbit          --------------------cgcgcgcggtcacgttttccactatgtgacagcggcgggt
Horse           --------------------cgcgcgcggtcacgttttccactatgtgacagcggcgagg
Sheep           -----------------------gagcggtcacgttttccactatgtgacagcggcgagt
Pika            ----------------------------------------actatgtgagagcgactaa-
                                                        ********* ****.  .

Mouse           ga--cgcggcggcagcggcgcta-----------ctgggactagcggctccgggcggc--
Rat             ga--cgcggtggcagcggcgcta-----------cagtgactagcggctccgggcggc--
GuineaPig       cg--ctcggcgcaggcggcgcgg-----------ggggaaccagctgctctaggtggc--
Human           cggccgcggcggaggcggcggcg------ctgaggggatacgtgcagctgtgggc-----
Rhesus          cggccgcggcggaggcggcggcg------ctgaggggctacgtgcagctgtgggcggtgg
Marmoset        cggcagcggcggaggcggcctcg------cggagggactgcgtgcggctgtgggcggt--
Rabbit          cgg-ggcggcggcggcggcgccg----------cgggctacgcgctggtctcggcggc--
Horse           cggcggcggcggcggcggctctg----------caggctacgcgctctcggcggtggt--
Sheep           ccgcggcggcggcggcggcagcgaacagccctgagggatacacgctgctttccgcggc--
Pika            ------------------------------------------------------------


Mouse           -------tgcggcg-------cagg-ccgagcgcaccaagtgacgggccgagcaagg---
Rat             -------tgcggcg-------cagg-ccgagcgcaccaagtgacgggtcgagcaaag---
GuineaPig       -------ggccgcgat-----cgtg-ccgggcggaccgaccgaggggccgactgaaggcc
Human           -------ggcggcggcgggcgcggggccgggcggacagagccgcgagtcggcggagg---
Rhesus          tggcggcggcggcagcgggcgcgggtccgggcggacagagccacgagtcggcggagg---
Marmoset        -------ggcggcggcaggcgcggggccgggcggacagagccgcgggccgacggagg---
Rabbit          -------ggcggcggcgggcgcggg-ccgggcgagcgcgcggactggccggcggagg---
Horse           -------ggcggccgggacaccgac-----------------------------gcg---
Sheep           -------ggcggcggcggggtcggc--------------------ggaccgcgcacg---
Pika            -------ggcagc--------cggt--------------------gaccggctgacggac
                        ** **        *.                               . *

Mouse           gacagacgcgcgggttgac------gcggcgaa-----gcg-cttattccagagcccgac
Rat             gaccgacgcgcgggtggac------gctgcgaa-----gcgcctcattccagagcccgac
GuineaPig       gacagacgcgaggccggacgcggcggcggtgagtgaacggaagctgctccag-------c
Human           gaccggcggacgggctgacgcgggcgcggcc-g-----gcgcttcgttccagagcccagc
Rhesus          gaccggcggacgggccgacgcgaacgcggccgg-----gcgcttcattccagagcccaac
Marmoset        gaccggcggactggccgacgcggacgcggccgg-----gcgcttcattccagagcccagc
Rabbit          gtccggcg-----------------acggctgg-----gcggttgactccggagcccacc
Horse           gacggccgggcgggcgggcgaggacgcggcgcg-----gcgcttcattcgggagcc-aac
Sheep           gatcgtccggcggactggttaggacgcggct-------ccggtctgccccggagc--agc
Pika            cgcggctgaggtgactgag------gcggctgagggctgaggttgattcgggagcccaac
                  . * .                  .* *.          .  . ...* .*       *
```

**Supplementary Figure 13 | Multiple sequence alignment of the *Per2* 5'UTR. (A)** Multiple sequence alignment of selected *Per2* 5'UTRs from different mammalian species using MAFFT (10). Identical nucleotides (asterisk) or pyrimidine or purine nucleotide conservation (period) among all species is shown. Note: only *Per2* transcriptional isoforms from each species containing the *Per2* uORF were used for the alignment.

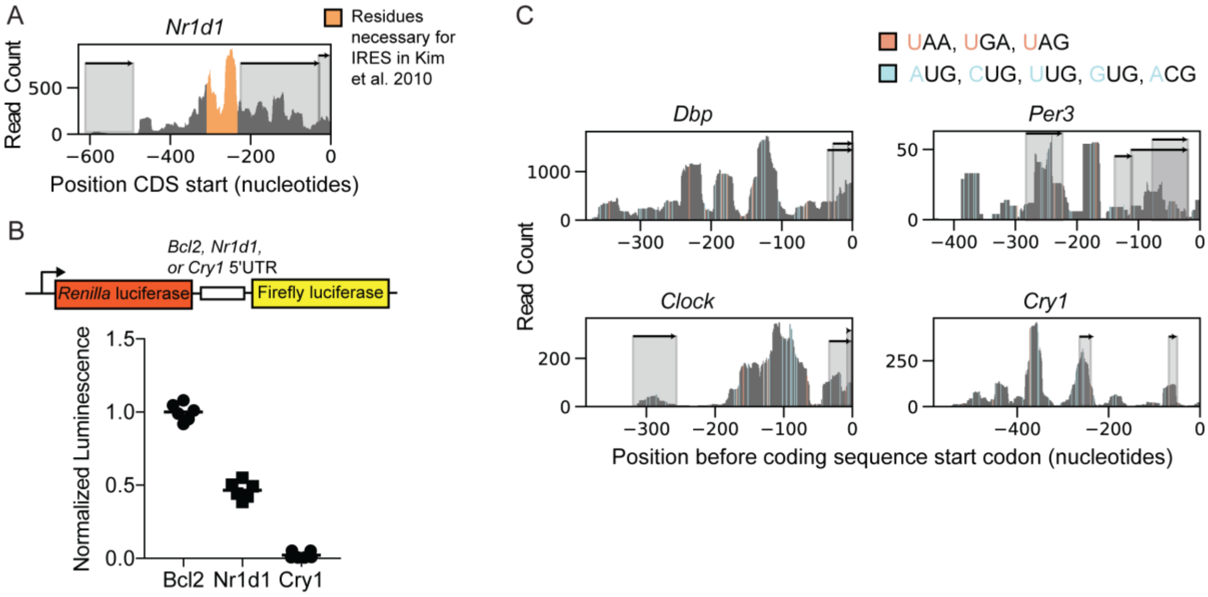


**Supplementary Figure 14 | Analysis of the IRES in the *Nrd1* 3'UTR and distribution of near-cognate start codons and stop codons in selected 5'UTRs. (A)** Extensive ribosome binding in *Nr1d1* 5' UTR overlaps with the region (orange) necessary for IRES-mediated translation of *Nr1d1* (11). **(B)** The efficiency of IRES-mediated translation for the *Nr1d1* 5'UTR and *Cry1* 5'UTR relative to a known IRES in *Bcl2* 5'UTR (12). **(C)** The 5' UTRs of *Dbp*, *Per3*, *Clock*, and *Cry1* have extensive ribosome binding in regions without a cognate uORF. Cognate and near-cognate start codons (light blue) and stop codons (orange) indicate regions of possible canonical uORFs (uORFs with an AUG start codon, shaded regions) and near-cognate uORFs (uORFs with a CUG, UUG, GUG, or ACG start codon).

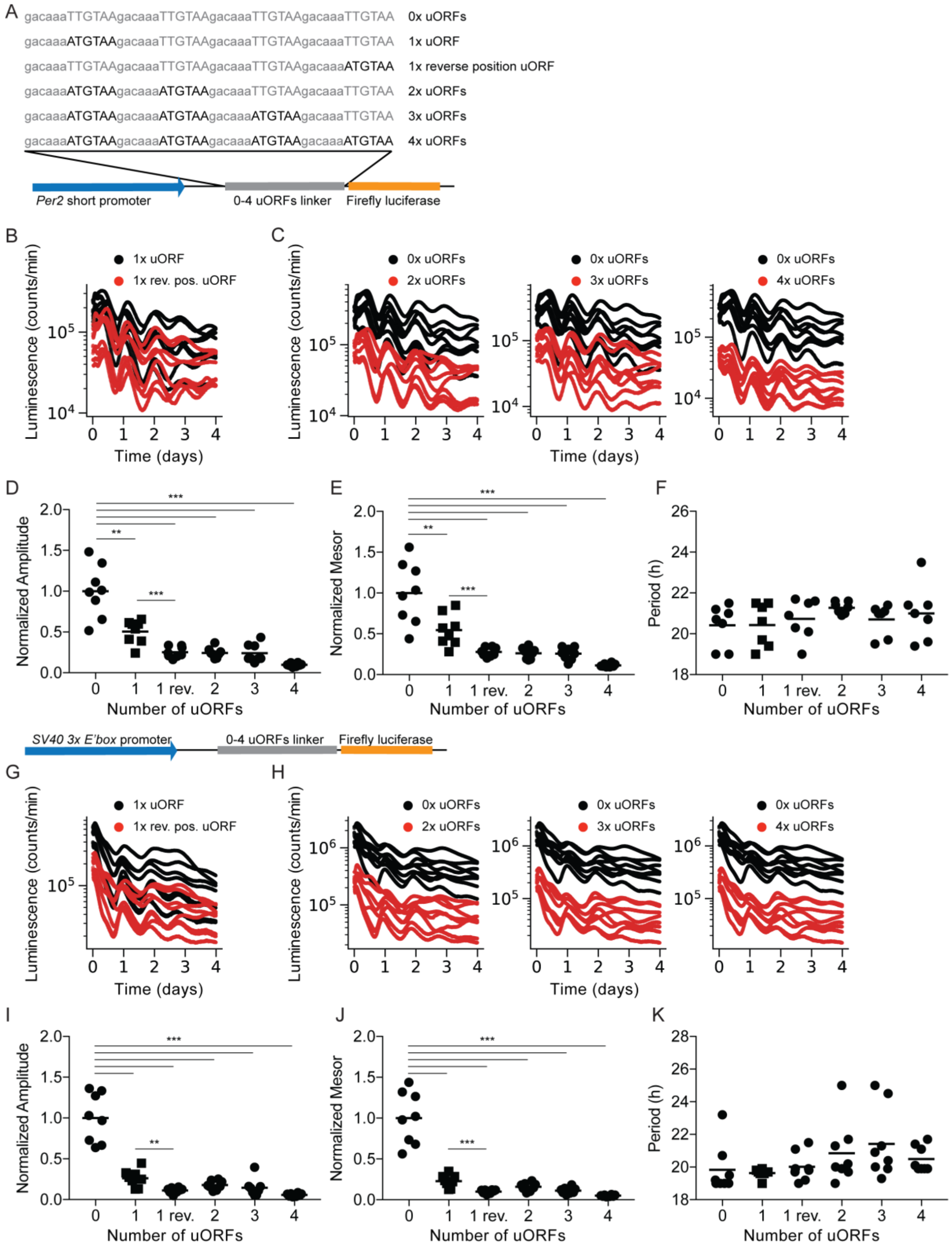


**Supplementary Figure 15 | Circadian rhythmicity analysis of reporter cells with variable numbers of uORFs. (A)** Construction of pGL3-P(*Per2*)-d*Luc* reporter plasmids with variable number of uORFs. **(B)** Bioluminescent recording of 3T3 cells transfected with the *Per2* short promoter containing exactly one uORF, 42 nt (1x uORF, black) or 6 nt (1x rev. pos. uORF, red) from the luciferase CDS. **(C)** Bioluminescent recording of 3T3 cells transfected with the *Per2* short promoter without an uORF (black) or containing (red) two uORFs (*left*), three uORFs (*middle*), or four uORFs (*right*). Cosinor analysis of the PMT traces in (**B**) and (**C**) showing amplitude (**D**), mesor (**E**), and period (**F**). **(G)** Bioluminescent recording of 3T3 cells transfected with the pGL3-P(SV40)-3xE'box-d*Luc* containing exactly one uORF, 42 nt (1x uORF, black) or 6 nt (1x rev. pos. uORF, red) from the luciferase CDS. **(H)** Bioluminescent recording of 3T3 cells transfected with pGL3-P(SV40)-3xE'box-d*Luc* without an uORF (black) or containing (red) two uORFs (*left*), three uORFs (*middle*), or four uORFs (*right*). Cosinor analysis of the PMT traces in (**G**) and (**H**) showing amplitude (**I**), mesor (**J**), and period (**K**).

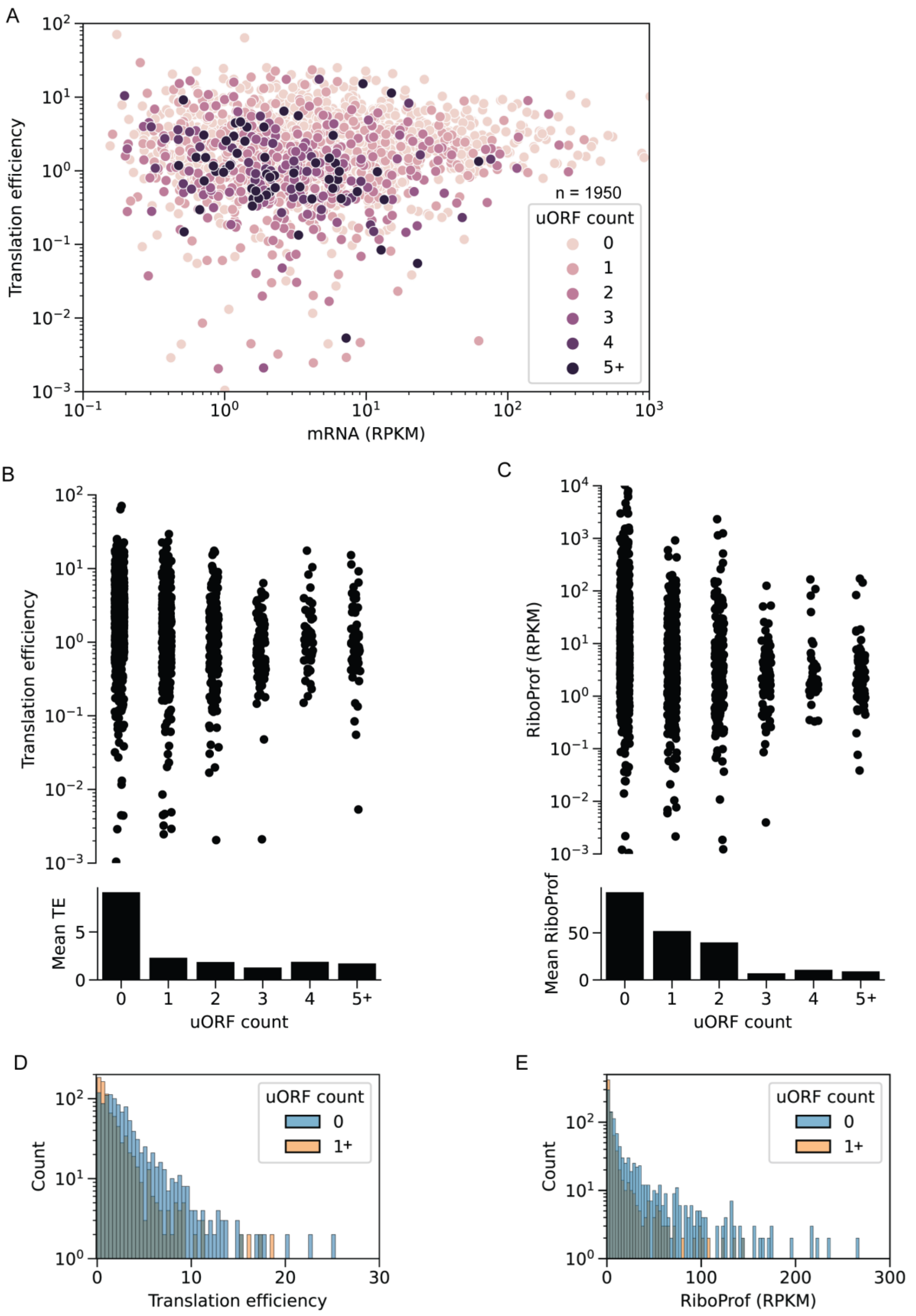


**Supplementary Figure 16 | Relationship between the number of uORFs and translation efficiency (TE) in the mouse liver using RNA-seq data from (13) and ribosome profiling**

**data from this study. (A)** Scatterplot of TE (mRNA RPKM from (13) divided by ribosome profiling RPKM) versus transcript abundance (RPKM) colored by the number of uORFs in each transcript. The distribution (*top*) and mean (*bottom*) of transcripts with the corresponding number of uORFs is shown for TE (**B**) and ribosome binding (**C**). Histograms of TE (**D**) and ribosome binding (**E**) for transcripts without an uORF (orange) or with one or more uORFs (blue).

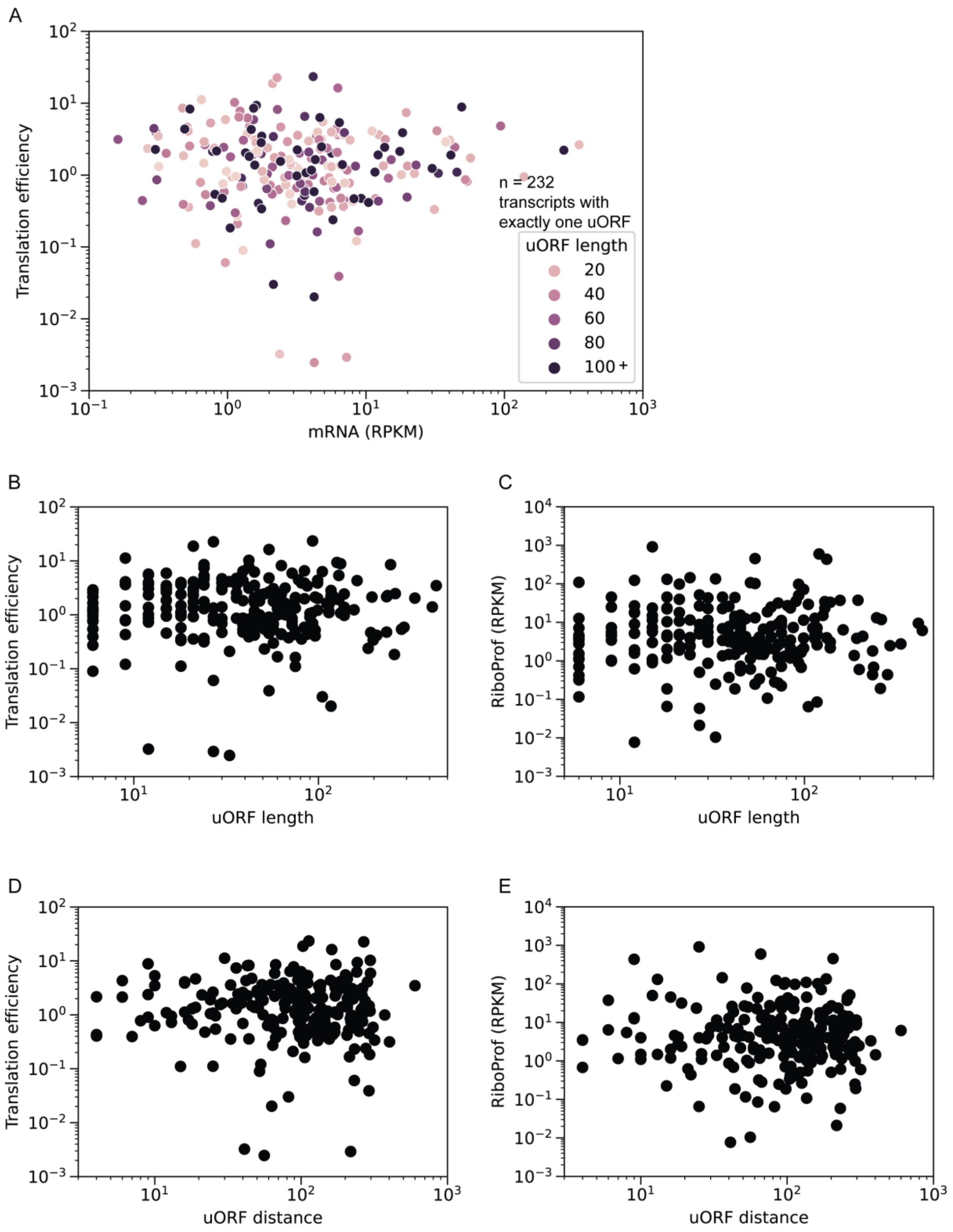


**Supplementary Figure 17 | Extended analysis of translation efficiency (TE) in the mouse liver using RNA-seq data from (13) and ribosome profiling data from this study. (A)** Scatterplot of TE (mRNA RPKM from (13) divided by ribosome profiling RPKM) versus transcript abundance (RPKM) colored by uORF length for each transcript with exactly one uORF. Scatterplot of TE (**B**) and ribosome binding (**C**) versus uORF length. Scatterplot of TE (**D**) and ribosome binding (**E**) versus distance of uORF to start of the CDS.

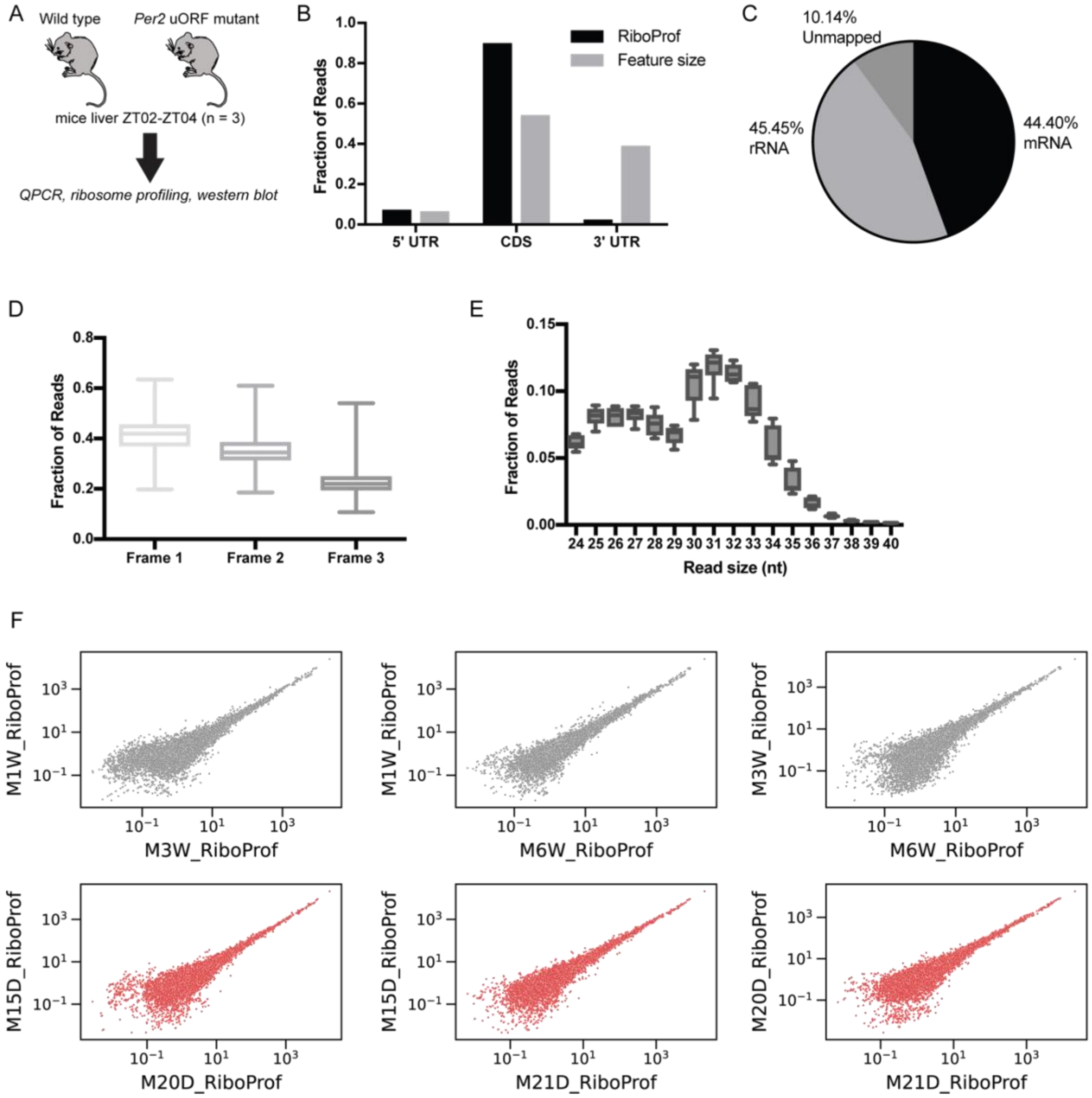


**Supplementary Figure 18 | Validation of ribosome profiling libraries from wild-type and *Per2* uORF mutant mice liver at ZT02-04 (related to Figure 4).** (**A**) Twelve to fourteen-week-old male wild-type and *Per2* uORF mutant mice under LD conditions were sacrificed at ZT02-ZT04 and liver sections were snap-frozen for ribosome profiling, total RNA sequencing, qPCR, and western blot analyses. (**B**) Read distribution within 5'UTRs, CDS, and 3'UTRs compared with the size of those features. Ribosome profiling reads are enriched for the CDS and 5'UTR, and relatively few reads map to the 3'UTR. (**C**) Frame analysis of ribosome profiling reads of single protein isoform mRNAs shows a preference for frame 1 (the coding frame). Box-and-whisker plots: midline, median; box, 25th and 75th percentiles. Whiskers are minimum and maximum values. (**D**) Box and whisker plots show the distribution of read lengths from all sequences. The majority of reads were between 27 and 33 nt, which matches the footprint size of the ribosome. (**E**) Summary of mapped reads.

(**F**) Pairwise correlation of ribosome profiling RPKM between wild-type replicates and between *Per2* uORF mutant replicates shows a high degree of correlation in ribosome binding between samples.

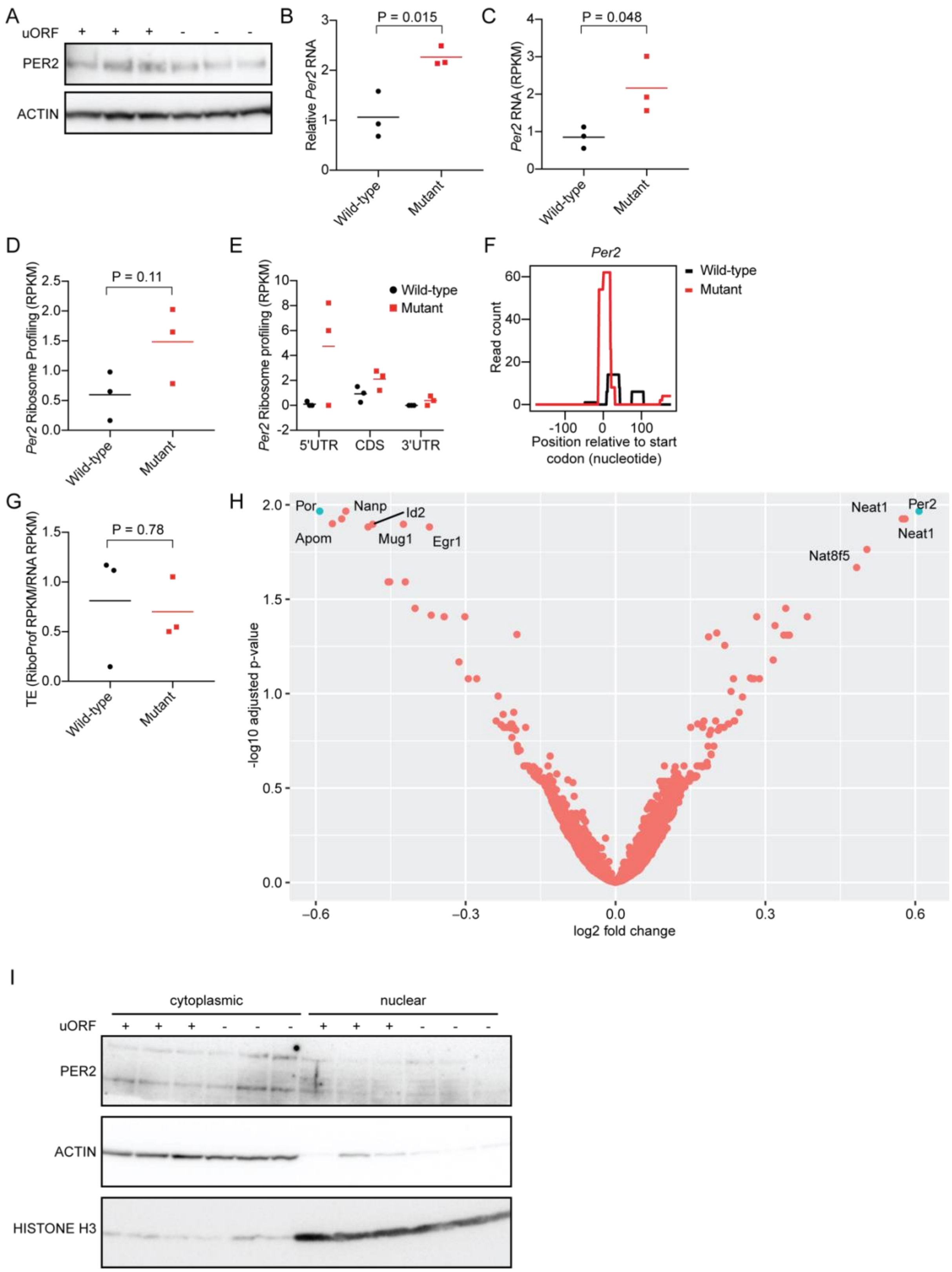


**Supplementary Figure 19 | Ribosome profiling, RNA sequencing, qPCR, and western blot analysis of liver from wild-type and *Per2* uORF mutant mice at ZT02-04 (related to Figure 4).** (**A**) Western blot of PER2 and actin protein of total protein lysate from wild-type (+ uORF) and *Per2* uORF mutant (- uORF) mice liver. Expression analysis of *Per2* mRNA from wild-type and *Per2* uORF mutant mice liver by qPCR (**B**) and total RNA sequencing (**C**). Ribosome binding (RPKM) across the entire *Per2* transcript **(D)** or compartmentalized by the 5'UTR, CDS, and 3'UTR (**E**). (**F**) Ribosome profiling reads in wild-type and *Per2* uORF mutant mice in the 5'UTR. Note the absence of reads across the uORF in both samples and an increased number of reads at the start codon in the *Per2* uORF mutant mice. (**G**) Translation efficiency (ribosome profiling RPKM divided by total RNA RPKM) in wild-type and *Per2* uORF mutant mice. (**H**) Differential expression analysis by DESeq2.0 (14) indicating genes with a significant change in gene expression as defined by an adjusted p-value < 0.05 and an absolute log2 fold change ≥ 0.58 (blue) shows that *Per2* mRNA is significantly increased in *Per2* uORF mutant mice. (**I**) Immunoblot analysis of PER2, actin, and histone H3 of cytoplasmic and nuclear lysates from wild-type (+ uORF) and *Per2* uORF mutant (- uORF) mice liver.

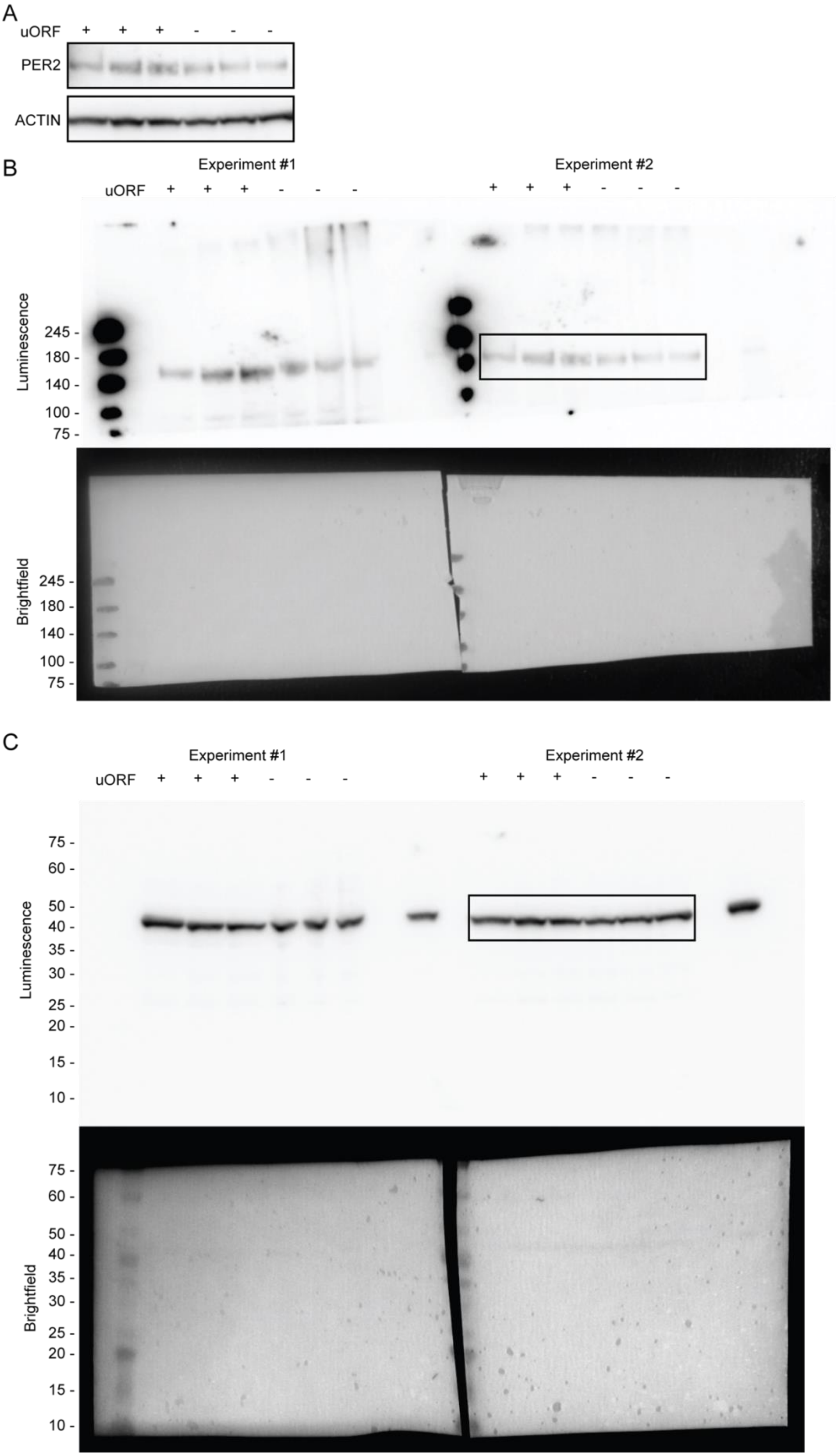
A
uORF + + + - - -
PER2
ACTIN
B
Experiment #1
Experiment #2
uORF + + + - - -
+ + + - - -
Luminescence
245 -
180 -
140 -
100 -
75 -
Brightfield
245 -
180 -
140 -
100 -
75 -
C
Experiment #1
Experiment #2
uORF + + + - - -
+ + + - - -
Luminescence
75 -
60 -
50 -
40 -
35 -
30 -
25 -
20 -
15 -
10 -
Brightfield
75 -
60 -
50 -
40 -
35 -
30 -
25 -
20 -
15 -
10 -

**Supplementary Figure 20 | Uncropped western blot images from Fig. S19A.** (**A**) Cropped western blot reproduced from Fig. S19A. Original uncropped images of the PER2 (**B**) and actin (**C**) western blot (*top*) and brightfield image showing the molecular weight ladder (*bottom*). Cropped region (black boxes).

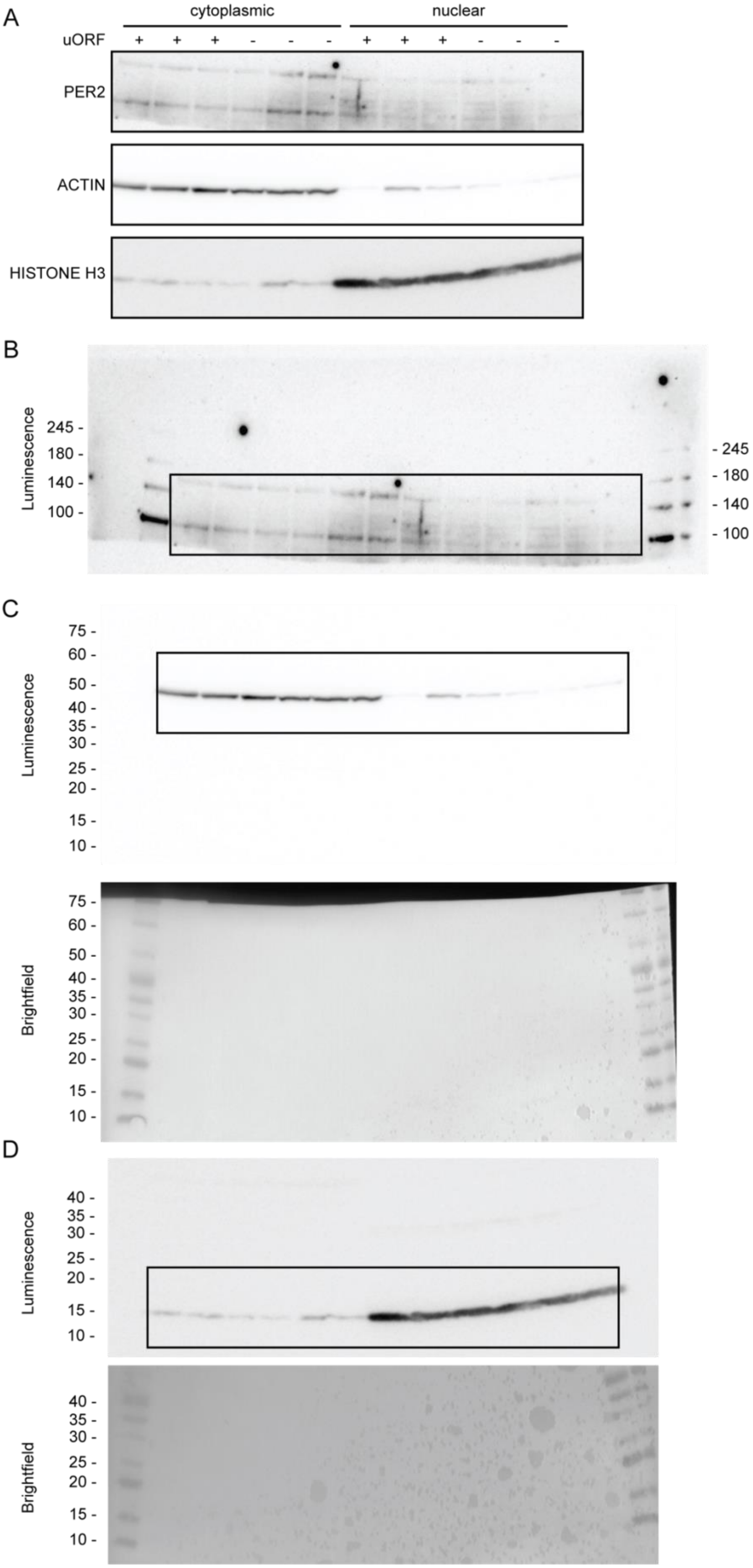
A
cytoplasmic
nuclear
uORF + + + - - - + + + - - -
PER2
ACTIN
HISTONE H3
B
Luminescence
245 -
180 -
140 -
100 -
- 245
- 180
- 140
- 100
C
Luminescence
75 -
60 -
50 -
40 -
35 -
30 -
25 -
20 -
15 -
10 -
Brightfield
75 -
60 -
50 -
40 -
35 -
30 -
25 -
20 -
15 -
10 -
D
Luminescence
40 -
35 -
30 -
25 -
20 -
15 -
10 -
Brightfield
40 -
35 -
30 -
25 -
20 -
15 -
10 -

**Supplementary Figure 21 | Uncropped western blot images from Fig. S19I.** (**A**) Cropped western blot reproduced from Fig. S19I. Original uncropped western blot of PER2 (**B**) with the molecular weight ladder visible in the outermost lanes. Uncropped images of actin (**C**) and histone H3 (**D**) western blot (*top*) and brightfield image showing the molecular weight ladder (*bottom*). Cropped region (black boxes).

**Table S1 | Sequence statistics for ribosome profiling data showing the circadian time (CT), number of reads per sample, number of reads mapped to rRNA, unmapped reads, and mapped reads.**

| Sample | CT | Reads | rRNA aligned | rRNA (%) | Unmapped | Unmapped (%) | Mapped | Mapped (%) |
|---|---|---|---|---|---|---|---|---|
| A1 | 0 | 67420894 | 31637068 | 47 | 6493665 | 10 | 29290161 | 43 |
| A2 | 4 | 79568956 | 40132748 | 50 | 6120169 | 8 | 33316039 | 42 |
| A3 | 8 | 92923717 | 46040656 | 50 | 7305061 | 8 | 39578000 | 43 |
| A4 | 12 | 89178791 | 37939665 | 43 | 4558237 | 5 | 46680889 | 52 |
| A5 | 16 | 73866781 | 37993296 | 51 | 3002837 | 4 | 32870648 | 44 |
| A6 | 20 | 57310601 | 29343482 | 51 | 3785209 | 7 | 24181910 | 42 |
| A7 | 24 | 57919810 | 30413247 | 53 | 1903635 | 3 | 25602928 | 44 |
| B1 | 0 | 82543812 | 46972438 | 57 | 5223484 | 6 | 30347890 | 37 |
| B2 | 4 | 76767842 | 45784936 | 60 | 2978293 | 4 | 28004613 | 36 |
| B3 | 8 | 67206206 | 31921429 | 47 | 2040538 | 3 | 33244239 | 49 |
| B4 | 12 | 77788251 | 44039811 | 57 | 2632701 | 3 | 31115739 | 40 |
| B5 | 16 | 73477498 | 42486921 | 58 | 2420565 | 3 | 28570012 | 39 |
| B6 | 20 | 79546292 | 41792698 | 53 | 4794899 | 6 | 32958695 | 41 |
| B7 | 24 | 91267817 | 37921444 | 42 | 20546228 | 23 | 32800145 | 36 |
| Total | | 1066787268 | 544419839 | 51 | 73805521 | 7 | 448561908 | 42 |

### Table S2 | JTK analysis of ribosome profiling data for circadian (BH.Q < 0.05) transcripts.

| Name | RefSeq ID | BH.Q | ADJ.P | PER | LAG | AMP | CT00A | CT00B | CT04A | CT04B | CT08A | CT08B | CT12A | CT12B | CT16A | CT16B | CT20A | CT20B | CT24A | CT24B |
|---|---|---|---|---|---|---|---|---|---|---|---|---|---|---|---|---|---|---|---|---|
| Sigmar1 | NM_001286541 | 0.00 | 1.23E-07 | 24 | 0 | 8.1 | 36.2 | 32.8 | 15.6 | 16.6 | 8.7 | 9.2 | 3.7 | 7.3 | 13.5 | 9.7 | 20.2 | 22.8 | 30.2 | 21.9 |
| Tardbp | NM_145556 | 0.01 | 5.99E-07 | 24 | 0 | 1.2 | 4.9 | 4.2 | 2.3 | 1.7 | 1.1 | 1.0 | 0.3 | 1.0 | 1.9 | 2.0 | 2.4 | 2.7 | 3.9 | 3.1 |
| Aqp8 | NM_001109045 | 0.01 | 2.20E-06 | 24 | 0 | 6.5 | 26.2 | 22.8 | 12.4 | 7.4 | 3.4 | 3.1 | 1.0 | 1.4 | 4.3 | 2.3 | 14.2 | 11.4 | 17.9 | 17.4 |
| Rdh11 | NM_021557 | 0.01 | 2.20E-06 | 24 | 0 | 5.6 | 27.3 | 21.1 | 12.2 | 9.5 | 7.5 | 9.4 | 4.1 | 3.3 | 9.5 | 7.6 | 14.7 | 20.0 | 19.6 | 16.1 |
| Arntl | NM_001243048 | 0.01 | 2.20E-06 | 24 | 0 | 2.2 | 4.7 | 4.1 | 2.0 | 2.4 | 0.0 | 0.3 | 0.0 | 0.1 | 1.5 | 1.3 | 2.6 | 3.8 | 3.7 | 3.0 |
| Ncald | NM_001170866 | 0.01 | 2.20E-06 | 24 | 4 | 1.7 | 5.0 | 4.6 | 5.7 | 5.5 | 3.5 | 3.4 | 2.1 | 1.1 | 2.4 | 2.0 | 3.3 | 3.4 | 5.1 | 5.1 |
| Ppard | NM_011145 | 0.01 | 2.20E-06 | 24 | 0 | 1.0 | 5.9 | 4.9 | 0.5 | 1.4 | 0.3 | 0.3 | 0.0 | 0.1 | 1.1 | 0.6 | 2.8 | 2.1 | 5.5 | 3.1 |
| Rnf144a | NM_001081977 | 0.01 | 2.20E-06 | 24 | 0 | 0.6 | 2.9 | 4.1 | 1.3 | 1.3 | 0.6 | 0.5 | 0.0 | 0.2 | 1.0 | 0.7 | 1.9 | 1.0 | 1.7 | 1.9 |
| Dennd2d | NM_001093754 | 0.01 | 2.20E-06 | 24 | 0 | 0.3 | 1.7 | 1.4 | 0.9 | 0.9 | 0.8 | 0.6 | 0.4 | 0.4 | 1.0 | 0.9 | 1.3 | 1.0 | 1.3 | 1.5 |
| Pnrc1 | NM_001033225 | 0.01 | 6.89E-06 | 24 | 0 | 25.7 | 59.5 | 68.6 | 43.2 | 40.4 | 16.3 | 11.6 | 11.1 | 8.5 | 34.7 | 16.5 | 34.4 | 44.1 | 44.1 | 49.7 |
| Aqp8 | NM_007474 | 0.01 | 6.89E-06 | 24 | 0 | 21.1 | 98.7 | 86.8 | 55.5 | 31.4 | 17.2 | 12.9 | 7.7 | 6.3 | 19.0 | 15.4 | 37.2 | 49.7 | 61.4 | 51.7 |
| Pgd | NM_001081274 | 0.01 | 6.89E-06 | 24 | 0 | 5.6 | 21.4 | 22.5 | 14.4 | 8.9 | 8.5 | 9.1 | 7.1 | 6.7 | 11.9 | 13.7 | 15.2 | 17.2 | 18.2 | 16.0 |
| Cks2 | NM_025415 | 0.01 | 6.89E-06 | 24 | 0 | 3.4 | 28.2 | 22.4 | 13.1 | 11.4 | 6.1 | 8.6 | 2.4 | 6.7 | 10.1 | 9.6 | 12.4 | 10.5 | 16.3 | 13.9 |
| Capn2 | NM_009794 | 0.01 | 6.89E-06 | 24 | 0 | 2.0 | 11.3 | 12.4 | 7.7 | 6.4 | 4.6 | 4.8 | 4.3 | 3.0 | 6.1 | 6.6 | 6.5 | 7.4 | 9.4 | 7.9 |
| Tjp2 | NM_001198985 | 0.01 | 6.89E-06 | 24 | 0 | 1.2 | 5.2 | 4.0 | 3.3 | 2.4 | 1.6 | 1.9 | 1.6 | 1.0 | 3.1 | 2.2 | 2.6 | 3.5 | 3.7 | 3.3 |
| Wee1 | NM_009516 | 0.01 | 6.89E-06 | 24 | 16 | 1.1 | 0.6 | 0.6 | 0.1 | 0.3 | 2.0 | 1.0 | 2.3 | 2.1 | 3.8 | 2.9 | 1.2 | 1.3 | 0.7 | 0.4 |
| Clpx | NM_001044389 | 0.02 | 1.89E-05 | 24 | 0 | 26.1 | 67.5 | 67.5 | 57.7 | 48.2 | 18.4 | 16.1 | 9.1 | 5.6 | 22.4 | 17.0 | 29.7 | 46.0 | 58.6 | 63.6 |
| Nampt | NM_021524 | 0.02 | 1.89E-05 | 24 | 18 | 11.4 | 14.6 | 13.2 | 6.6 | 6.9 | 7.4 | 9.3 | 31.1 | 22.1 | 43.9 | 32.4 | 18.2 | 20.5 | 13.3 | 10.9 |
| Pmvk | NM_026784 | 0.02 | 1.89E-05 | 24 | 0 | 6.8 | 40.6 | 32.2 | 9.7 | 7.2 | 4.6 | 5.3 | 4.2 | 4.3 | 8.9 | 9.5 | 18.5 | 24.3 | 23.0 | 16.8 |
| Nat8 | NM_023455 | 0.02 | 1.89E-05 | 24 | 22 | 4.2 | 7.1 | 9.3 | 5.8 | 4.3 | 2.7 | 3.3 | 3.3 | 3.1 | 8.4 | 7.6 | 10.2 | 9.8 | 8.8 | 8.5 |
| Evi5 | NM_007964 | 0.02 | 1.89E-05 | 24 | 0 | 2.5 | 11.3 | 13.3 | 9.8 | 8.5 | 5.6 | 5.5 | 5.1 | 2.6 | 7.9 | 6.0 | 6.7 | 8.9 | 9.1 | 9.9 |
| Dpp9 | NM_172624 | 0.02 | 1.89E-05 | 24 | 0 | 2.0 | 12.0 | 10.9 | 7.5 | 6.9 | 4.7 | 4.0 | 2.4 | 3.4 | 5.9 | 4.9 | 6.2 | 6.6 | 10.5 | 7.1 |
| Tjp3 | NM_001282095 | 0.02 | 1.89E-05 | 24 | 0 | 1.8 | 5.5 | 5.4 | 3.8 | 2.0 | 1.0 | 1.3 | 0.7 | 0.6 | 3.5 | 1.8 | 3.2 | 3.2 | 4.4 | 4.0 |
| Bloc1s5 | NM_139063 | 0.02 | 1.89E-05 | 24 | 0 | 1.6 | 13.6 | 11.4 | 8.2 | 7.8 | 6.6 | 5.5 | 4.0 | 5.5 | 7.8 | 6.5 | 7.0 | 8.8 | 10.0 | 8.6 |
| Gpcpd1 | NM_001042672 | 0.02 | 1.89E-05 | 24 | 12 | 1.5 | 3.0 | 2.4 | 3.8 | 3.2 | 9.1 | 4.8 | 23.0 | 11.7 | 4.5 | 4.3 | 3.0 | 2.9 | 2.1 | 1.9 |
| Dock6 | NM_177030 | 0.02 | 1.89E-05 | 24 | 0 | 0.8 | 3.2 | 2.7 | 1.9 | 1.2 | 0.8 | 0.8 | 0.5 | 0.4 | 1.4 | 1.4 | 1.6 | 2.0 | 2.3 | 1.7 |
| Fam222a | NM_001004180 | 0.02 | 1.89E-05 | 24 | 0 | 0.5 | 2.4 | 3.4 | 0.4 | 1.1 | 0.7 | 0.4 | 0.0 | 0.2 | 0.9 | 0.7 | 1.2 | 1.4 | 2.2 | 1.2 |
| Elovl3 | NM_007703 | 0.03 | 4.69E-05 | 24 | 1 | 182.4 | 634.0 | 590.9 | 326.6 | 365.0 | 49.3 | 64.1 | 27.1 | 24.1 | 99.2 | 40.9 | 209.9 | 318.8 | 529.0 | 450.4 |
| Hacl1 | NM_019975 | 0.03 | 4.69E-05 | 24 | 0 | 77.4 | 247.4 | 188.7 | 134.6 | 114.1 | 75.0 | 64.3 | 46.6 | 42.3 | 140.7 | 84.9 | 147.7 | 199.3 | 194.5 | 180.9 |
| Cyp8b1 | NM_010012 | 0.03 | 4.69E-05 | 24 | 2 | 60.6 | 250.8 | 224.1 | 172.1 | 160.4 | 92.1 | 75.7 | 47.1 | 51.5 | 74.6 | 80.9 | 122.2 | 134.5 | 207.2 | 165.3 |
| Adck3 | NM_001163290 | 0.03 | 4.69E-05 | 24 | 0 | 44.8 | 192.7 | 178.5 | 142.5 | 97.0 | 66.5 | 36.9 | 24.3 | 21.4 | 70.3 | 44.7 | 69.5 | 100.3 | 143.2 | 123.5 |
| Pnp | NM_013632 | 0.03 | 4.69E-05 | 24 | 0 | 11.6 | 39.3 | 43.4 | 33.2 | 29.3 | 14.7 | 16.0 | 13.6 | 8.5 | 26.6 | 20.4 | 18.6 | 27.3 | 33.3 | 34.5 |
| Dhcr7 | NM_007856 | 0.03 | 4.69E-05 | 24 | 0 | 10.3 | 75.9 | 59.3 | 33.6 | 30.7 | 23.2 | 29.5 | 24.1 | 13.9 | 25.5 | 34.6 | 33.5 | 49.8 | 52.2 | 52.5 |
| Efna1 | NM_001162425 | 0.03 | 4.69E-05 | 24 | 0 | 8.6 | 44.0 | 29.9 | 25.3 | 25.9 | 17.4 | 19.4 | 9.0 | 12.3 | 24.4 | 15.1 | 31.6 | 27.2 | 41.9 | 26.6 |
| Nr1d1 | NM_145434 | 0.03 | 4.69E-05 | 24 | 10 | 7.0 | 1.3 | 1.3 | 20.9 | 8.1 | 32.1 | 18.8 | 6.8 | 9.2 | 5.6 | 2.0 | 0.7 | 0.8 | 1.5 | 0.8 |
| Caprin1 | NM_001111290 | 0.03 | 4.69E-05 | 24 | 0 | 6.3 | 30.6 | 24.1 | 19.0 | 13.9 | 10.1 | 9.5 | 7.6 | 5.9 | 18.6 | 15.5 | 17.9 | 18.0 | 22.7 | 22.9 |
| Gale | NM_178389 | 0.03 | 4.69E-05 | 24 | 0 | 5.1 | 21.9 | 16.3 | 9.8 | 8.2 | 2.8 | 5.1 | 1.9 | 2.2 | 8.2 | 10.9 | 12.4 | 11.0 | 12.1 | 10.9 |
| Avpr1a | NM_016847 | 0.03 | 4.69E-05 | 24 | 0 | 5.0 | 22.8 | 20.0 | 12.6 | 10.7 | 7.8 | 5.0 | 3.0 | 1.6 | 9.8 | 6.2 | 7.3 | 12.1 | 16.3 | 14.8 |
| Asb13 | NM_178283 | 0.03 | 4.69E-05 | 24 | 0 | 4.7 | 20.5 | 20.2 | 14.3 | 11.4 | 7.7 | 7.4 | 5.5 | 5.7 | 12.3 | 8.4 | 10.9 | 12.6 | 16.1 | 12.9 |
| Mmab | NM_029956 | 0.03 | 4.69E-05 | 24 | 0 | 3.0 | 11.5 | 10.0 | 7.5 | 5.1 | 5.1 | 3.5 | 3.0 | 3.0 | 5.2 | 4.3 | 7.8 | 7.3 | 5.6 | 9.1 |
| Rfxank | NM_001025589 | 0.03 | 4.69E-05 | 24 | 0 | 2.7 | 9.0 | 7.4 | 5.7 | 6.7 | 2.2 | 2.5 | 2.4 | 1.9 | 5.7 | 3.6 | 6.7 | 6.4 | 9.3 | 7.8 |
| Lgalsl | NM_173752 | 0.03 | 4.69E-05 | 24 | 0 | 2.6 | 10.8 | 6.4 | 5.0 | 4.4 | 1.5 | 1.2 | 0.7 | 0.5 | 5.5 | 3.2 | 5.2 | 6.7 | 7.2 | 6.7 |
| Chka | NM_013490 | 0.03 | 4.69E-05 | 24 | 0 | 2.5 | 7.7 | 6.3 | 4.8 | 5.0 | 1.3 | 1.5 | 0.2 | 0.5 | 3.6 | 2.0 | 2.3 | 4.6 | 6.3 | 5.8 |
| Camk1d | NM_177343 | 0.03 | 4.69E-05 | 24 | 2 | 1.2 | 6.6 | 4.2 | 3.8 | 3.3 | 2.3 | 2.6 | 2.5 | 0.9 | 0.8 | 1.2 | 2.9 | 2.8 | 6.1 | 5.1 |
| Per2 | NM_011066 | 0.03 | 4.69E-05 | 24 | 18 | 1.2 | 2.8 | 1.5 | 0.7 | 0.2 | 1.3 | 1.2 | 2.0 | 2.0 | 6.6 | 4.8 | 3.2 | 2.8 | 2.1 | 1.6 |
| St5 | NM_001001326 | 0.03 | 4.69E-05 | 24 | 0 | 0.8 | 2.1 | 2.6 | 1.9 | 1.4 | 0.8 | 0.7 | 0.3 | 0.3 | 1.3 | 1.0 | 1.4 | 1.9 | 2.2 | 1.5 |
| Dus2 | NM_025518 | 0.03 | 4.69E-05 | 24 | 2 | 0.6 | 3.5 | 2.6 | 2.5 | 2.5 | 0.7 | 1.8 | 1.5 | 1.4 | 2.0 | 1.8 | 2.2 | 2.2 | 3.8 | 3.4 |
| Slc34a2 | NM_011402 | 0.03 | 4.69E-05 | 24 | 0 | 0.6 | 5.4 | 3.0 | 0.6 | 0.8 | 0.0 | 0.2 | 0.0 | 0.0 | 1.1 | 0.2 | 0.4 | 1.3 | 2.3 | 2.6 |
| Ccdc151 | NM_001163787 | 0.03 | 4.69E-05 | 24 | 20 | 0.3 | 0.7 | 0.7 | 0.6 | 0.3 | 0.1 | 0.3 | 0.7 | 0.3 | 1.6 | 0.8 | 0.7 | 1.0 | 0.7 | 0.7 |
| Cyb5b | NM_025558 | 0.03 | 1.07E-04 | 24 | 0 | 50.9 | 283.6 | 239.2 | 169.2 | 130.8 | 106.1 | 82.4 | 75.6 | 67.5 | 149.0 | 97.2 | 126.9 | 148.2 | 194.2 | 173.6 |
| Eif4ebp3 | NM_201256 | 0.03 | 1.07E-04 | 24 | 18 | 43.8 | 97.9 | 75.0 | 30.9 | 24.6 | 53.1 | 18.2 | 187.2 | 100.8 | 396.5 | 111.1 | 92.3 | 110.7 | 56.6 | 64.6 |

| | | | | | | | | | | | | | | | | | | | | |
|---|---|---|---|---|---|---|---|---|---|---|---|---|---|---|---|---|---|---|---|---|
| Gys2 | NM_145572 | 0.03 | 1.07E-04 | 24 | 16 | 22.4 | 45.8 | 41.6 | 31.0 | 21.7 | 58.8 | 36.8 | 85.5 | 70.9 | 103.3 | 86.3 | 40.5 | 53.7 | 39.0 | 36.6 |
| Nop10 | NM_025403 | 0.03 | 1.07E-04 | 24 | 0 | 19.1 | 197.0 | 170.9 | 123.8 | 122.5 | 107.9 | 102.6 | 86.0 | 95.2 | 188.9 | 115.7 | 129.7 | 135.0 | 147.9 | 135.6 |
| Mrap | NM_029844 | 0.03 | 1.07E-04 | 24 | 0 | 13.4 | 54.0 | 47.0 | 33.6 | 24.9 | 16.4 | 14.5 | 12.6 | 12.1 | 35.3 | 19.3 | 35.4 | 28.4 | 33.3 | 41.7 |
| Acss2 | NM_019811 | 0.03 | 1.07E-04 | 24 | 0 | 12.6 | 80.3 | 62.4 | 36.2 | 21.1 | 23.1 | 16.3 | 18.3 | 13.5 | 26.8 | 25.7 | 30.0 | 58.4 | 53.5 | 48.8 |
| Ppp1r3c | NM_016854 | 0.03 | 1.07E-04 | 24 | 1 | 10.7 | 77.1 | 39.5 | 25.9 | 24.0 | 12.0 | 11.7 | 11.1 | 4.3 | 18.4 | 9.5 | 19.5 | 38.5 | 62.1 | 56.4 |
| Sgpl1 | NM_009163 | 0.03 | 1.07E-04 | 24 | 0 | 7.1 | 30.3 | 22.3 | 21.8 | 15.8 | 13.2 | 11.6 | 9.9 | 6.8 | 19.6 | 17.6 | 15.9 | 21.8 | 24.8 | 25.0 |
| Hypk | NM_026318 | 0.03 | 1.07E-04 | 24 | 0 | 7.0 | 62.3 | 58.0 | 28.7 | 33.7 | 31.7 | 27.0 | 19.0 | 22.6 | 34.5 | 33.4 | 42.5 | 36.9 | 39.9 | 35.7 |
| Pter | NM_008961 | 0.03 | 1.07E-04 | 24 | 2 | 5.8 | 28.1 | 24.0 | 22.0 | 17.4 | 13.2 | 15.0 | 10.6 | 7.3 | 16.9 | 12.5 | 15.9 | 20.7 | 28.6 | 21.5 |
| Tfpi2 | NM_009364 | 0.03 | 1.07E-04 | 24 | 0 | 5.8 | 34.1 | 31.4 | 21.1 | 20.7 | 13.0 | 16.8 | 12.2 | 8.3 | 17.0 | 14.4 | 12.3 | 25.8 | 27.0 | 29.7 |
| Slc16a1 | NM_009196 | 0.03 | 1.07E-04 | 24 | 0 | 5.5 | 33.5 | 25.6 | 16.4 | 14.3 | 10.0 | 11.4 | 8.3 | 8.2 | 20.8 | 14.6 | 14.5 | 20.8 | 21.6 | 21.1 |
| Tgm2 | NM_009373 | 0.03 | 1.07E-04 | 24 | 0 | 5.4 | 58.5 | 37.5 | 25.8 | 20.6 | 18.1 | 16.3 | 13.1 | 10.9 | 24.7 | 24.1 | 24.2 | 23.2 | 35.4 | 27.5 |
| Btg1 | NM_007569 | 0.03 | 1.07E-04 | 24 | 0 | 5.1 | 16.9 | 16.3 | 11.4 | 7.9 | 6.0 | 5.1 | 3.6 | 2.8 | 9.5 | 3.3 | 6.3 | 12.3 | 12.6 | 13.5 |
| Cdkn1a | NM_001111099 | 0.03 | 1.07E-04 | 24 | 0 | 3.9 | 22.0 | 14.0 | 4.5 | 5.7 | 0.4 | 2.3 | 0.5 | 0.6 | 6.6 | 2.6 | 7.8 | 5.7 | 7.6 | 12.4 |
| Spryd7 | NM_025697 | 0.03 | 1.07E-04 | 24 | 0 | 3.7 | 20.5 | 16.4 | 13.5 | 9.2 | 8.6 | 8.2 | 7.2 | 7.5 | 14.5 | 9.2 | 15.0 | 12.4 | 14.7 | 13.8 |
| Acacb | NM_133904 | 0.03 | 1.07E-04 | 24 | 0 | 3.7 | 17.5 | 13.0 | 6.8 | 4.8 | 2.8 | 2.7 | 3.9 | 2.7 | 5.9 | 4.7 | 8.5 | 13.7 | 12.7 | 10.7 |
| Pi4k2a | NM_145501 | 0.03 | 1.07E-04 | 24 | 0 | 3.4 | 17.6 | 16.8 | 14.7 | 10.8 | 8.4 | 8.1 | 5.6 | 7.3 | 13.7 | 9.4 | 10.5 | 12.0 | 15.6 | 14.0 |
| Lsr | NM_001164185 | 0.03 | 1.07E-04 | 24 | 0 | 2.8 | 14.7 | 11.8 | 5.8 | 7.9 | 4.8 | 5.3 | 4.2 | 4.7 | 10.8 | 6.2 | 9.5 | 8.1 | 11.9 | 10.2 |
| Ddx17 | NM_001040187 | 0.03 | 1.07E-04 | 24 | 0 | 2.3 | 17.6 | 10.2 | 8.4 | 6.9 | 4.8 | 5.1 | 4.5 | 3.4 | 7.8 | 7.0 | 7.4 | 8.2 | 13.3 | 11.5 |
| Ext2 | NM_010163 | 0.03 | 1.07E-04 | 24 | 0 | 1.5 | 9.1 | 10.2 | 5.9 | 4.8 | 4.6 | 3.7 | 3.1 | 3.0 | 5.9 | 3.4 | 5.8 | 5.2 | 6.6 | 7.0 |
| Ncoa5 | NM_144892 | 0.03 | 1.07E-04 | 24 | 0 | 1.0 | 5.3 | 3.9 | 2.3 | 1.9 | 1.4 | 1.3 | 0.9 | 1.0 | 3.5 | 2.0 | 2.1 | 2.8 | 3.6 | 4.1 |
| Atg7 | NM_001253717 | 0.03 | 1.07E-04 | 24 | 0 | 0.9 | 6.0 | 5.5 | 4.1 | 3.4 | 2.8 | 2.5 | 2.1 | 2.2 | 4.0 | 3.4 | 3.5 | 3.1 | 4.8 | 4.4 |
| Slc17a8 | NM_182959 | 0.03 | 1.07E-04 | 24 | 0 | 0.9 | 3.3 | 3.1 | 2.2 | 1.1 | 0.9 | 0.5 | 0.3 | 0.9 | 1.3 | 1.4 | 2.2 | 1.7 | 2.3 | 1.7 |
| Agpat5 | NM_026792 | 0.03 | 1.07E-04 | 24 | 0 | 0.9 | 6.0 | 7.4 | 3.7 | 4.2 | 3.0 | 3.4 | 1.4 | 2.5 | 5.1 | 4.1 | 4.0 | 4.6 | 5.4 | 5.7 |
| Rbmx | NM_001166623 | 0.03 | 1.07E-04 | 24 | 0 | 0.8 | 4.0 | 2.9 | 2.3 | 1.5 | 1.4 | 1.5 | 0.7 | 1.1 | 2.2 | 1.1 | 2.3 | 2.7 | 3.2 | 2.5 |
| Dut | NM_023595 | 0.03 | 1.07E-04 | 24 | 0 | 0.5 | 5.5 | 4.2 | 2.2 | 2.5 | 2.6 | 2.0 | 0.7 | 1.1 | 2.0 | 2.0 | 5.4 | 2.6 | 3.3 | 3.1 |
| Mthfr | NM_001161798 | 0.03 | 1.07E-04 | 24 | 2 | 0.4 | 1.6 | 1.0 | 1.0 | 0.9 | 0.2 | 0.2 | 0.0 | 0.1 | 0.8 | 0.4 | 0.5 | 0.8 | 1.6 | 1.2 |
| Per3 | NM_011067 | 0.03 | 1.07E-04 | 24 | 16 | 0.4 | 0.1 | 0.2 | 0.0 | 0.0 | 0.5 | 0.4 | 1.4 | 0.8 | 1.0 | 0.9 | 0.5 | 0.3 | 0.2 | 0.2 |
| Oma1 | NM_025909 | 0.03 | 1.07E-04 | 24 | 0 | 0.3 | 8.6 | 7.6 | 5.2 | 5.1 | 5.1 | 4.9 | 3.3 | 2.6 | 6.6 | 4.9 | 5.4 | 6.0 | 6.4 | 6.0 |
| Mtrf1 | NM_145960 | 0.03 | 1.07E-04 | 24 | 0 | 0.3 | 2.8 | 2.4 | 2.0 | 2.1 | 1.4 | 1.2 | 1.9 | 1.2 | 1.9 | 1.7 | 2.1 | 2.0 | 2.3 | 2.5 |
| St5 | NM_029811 | 0.03 | 1.07E-04 | 24 | 2 | 0.2 | 0.3 | 0.4 | 0.1 | 0.2 | 0.1 | 0.1 | 0.0 | 0.0 | 0.1 | 0.0 | 0.4 | 0.2 | 0.4 | 0.4 |
| Plekhg1 | NM_001033253 | 0.03 | 1.07E-04 | 24 | 0 | 0.1 | 0.8 | 0.8 | 0.5 | 0.6 | 0.4 | 0.3 | 0.5 | 0.3 | 0.5 | 0.5 | 0.6 | 0.6 | 0.7 | 0.7 |
| Eif2ak4 | NM_013719 | 0.03 | 1.07E-04 | 24 | 0 | 0.1 | 0.5 | 0.6 | 0.2 | 0.2 | 0.1 | 0.1 | 0.1 | 0.1 | 0.3 | 0.2 | 0.5 | 0.3 | 0.4 | 0.3 |
| Creb3l1 | NM_011957 | 0.03 | 1.07E-04 | 24 | 0 | 0.1 | 0.4 | 0.6 | 0.2 | 0.2 | 0.1 | 0.1 | 0.1 | 0.1 | 0.3 | 0.2 | 0.2 | 0.3 | 0.3 | 0.3 |
| Prdm10 | NM_001080817 | 0.03 | 1.07E-04 | 24 | 0 | 0.1 | 0.5 | 0.6 | 0.3 | 0.4 | 0.4 | 0.3 | 0.2 | 0.2 | 0.3 | 0.3 | 0.4 | 0.4 | 0.5 | 0.4 |
| Bhmt | NM_016668 | 0.05 | 2.29E-04 | 24 | 0 | 57.1 | 362.2 | 255.4 | 289.4 | 185.9 | 124.4 | 119.1 | 114.4 | 86.9 | 175.5 | 136.1 | 138.8 | 248.3 | 245.3 | 318.2 |
| Plin2 | NM_007408 | 0.05 | 2.29E-04 | 24 | 2 | 46.8 | 236.4 | 188.9 | 145.4 | 159.6 | 137.3 | 93.5 | 56.2 | 66.7 | 118.5 | 90.0 | 106.9 | 141.7 | 195.0 | 145.1 |
| S100a10 | NM_009112 | 0.05 | 2.29E-04 | 24 | 0 | 42.9 | 251.9 | 240.6 | 210.9 | 169.6 | 110.4 | 111.9 | 94.9 | 66.3 | 153.0 | 110.9 | 128.9 | 160.6 | 191.6 | 193.6 |
| Cyb5r3 | NM_029787 | 0.05 | 2.29E-04 | 24 | 0 | 35.4 | 317.2 | 278.3 | 210.4 | 187.7 | 184.2 | 160.3 | 157.6 | 146.6 | 258.5 | 188.5 | 190.8 | 233.0 | 253.0 | 235.9 |
| Sdc1 | NM_011519 | 0.05 | 2.29E-04 | 24 | 0 | 33.1 | 203.0 | 165.9 | 96.1 | 85.7 | 60.3 | 47.8 | 63.9 | 46.1 | 144.9 | 71.1 | 107.8 | 105.6 | 129.4 | 117.2 |
| Tubb2a | NM_009450 | 0.05 | 2.29E-04 | 24 | 0 | 29.5 | 79.5 | 60.2 | 51.5 | 36.8 | 5.7 | 9.8 | 4.8 | 2.4 | 37.9 | 17.9 | 27.2 | 39.2 | 83.6 | 51.2 |
| Ddc | NM_001190448 | 0.05 | 2.29E-04 | 24 | 0 | 19.3 | 81.0 | 59.3 | 45.5 | 47.0 | 18.4 | 22.2 | 13.2 | 12.9 | 54.8 | 37.3 | 43.1 | 56.8 | 52.4 | 55.4 |
| Gclc | NM_010295 | 0.05 | 2.29E-04 | 24 | 2 | 16.2 | 66.9 | 71.1 | 60.3 | 51.2 | 44.5 | 29.1 | 26.7 | 23.5 | 35.5 | 28.3 | 31.7 | 44.7 | 63.3 | 54.3 |
| Tars | NM_033074 | 0.05 | 2.29E-04 | 24 | 0 | 15.6 | 62.1 | 52.9 | 43.9 | 33.1 | 20.1 | 18.8 | 14.3 | 12.7 | 35.3 | 27.6 | 26.4 | 40.9 | 49.9 | 40.4 |
| Pnkd | NM_001039509 | 0.05 | 2.29E-04 | 24 | 0 | 12.5 | 46.8 | 41.1 | 32.1 | 22.8 | 14.5 | 9.8 | 9.1 | 6.7 | 30.0 | 14.6 | 21.8 | 25.1 | 35.1 | 26.7 |
| Hmgcs1 | NM_145942 | 0.05 | 2.29E-04 | 24 | 0 | 11.2 | 62.0 | 58.9 | 37.6 | 26.4 | 23.5 | 41.8 | 16.9 | 10.5 | 25.0 | 21.7 | 31.3 | 56.0 | 57.4 | 54.0 |
| Cldn2 | NM_016675 | 0.05 | 2.29E-04 | 24 | 0 | 11.0 | 49.0 | 42.0 | 35.3 | 31.6 | 20.9 | 19.7 | 14.2 | 16.8 | 32.8 | 20.6 | 19.7 | 37.2 | 38.8 | 35.5 |
| Acsl4 | NM_001033600 | 0.05 | 2.29E-04 | 24 | 20 | 10.4 | 31.2 | 26.2 | 18.7 | 17.0 | 14.2 | 16.6 | 23.7 | 16.2 | 31.7 | 35.1 | 30.2 | 38.0 | 29.8 | 27.1 |
| Adh4 | NM_011996 | 0.05 | 2.29E-04 | 24 | 2 | 9.8 | 57.5 | 58.3 | 51.4 | 43.1 | 43.5 | 42.5 | 32.3 | 22.4 | 42.4 | 37.5 | 30.9 | 51.7 | 58.5 | 64.0 |
| Ran | NM_009391 | 0.05 | 2.29E-04 | 24 | 0 | 8.6 | 63.1 | 52.5 | 34.4 | 25.9 | 21.7 | 23.6 | 20.5 | 21.5 | 36.9 | 28.7 | 26.9 | 33.8 | 50.7 | 38.4 |
| Fdft1 | NM_010191 | 0.05 | 2.29E-04 | 24 | 0 | 6.2 | 22.9 | 21.4 | 7.1 | 7.5 | 6.6 | 13.6 | 6.4 | 4.2 | 10.7 | 11.3 | 15.4 | 17.9 | 19.6 | 18.6 |
| Mal2 | NM_178920 | 0.05 | 2.29E-04 | 24 | 0 | 5.7 | 24.9 | 24.0 | 11.4 | 9.4 | 5.8 | 4.8 | 4.2 | 6.9 | 22.2 | 9.2 | 14.9 | 13.8 | 17.8 | 18.0 |
| Atg101 | NM_026566 | 0.05 | 2.29E-04 | 24 | 0 | 5.4 | 28.0 | 24.8 | 20.2 | 16.7 | 12.8 | 11.9 | 7.2 | 8.3 | 18.9 | 11.0 | 13.9 | 17.0 | 22.8 | 21.1 |
| 2610528J11Rik | NM_025572 | 0.05 | 2.29E-04 | 24 | 0 | 4.9 | 16.4 | 16.6 | 13.3 | 9.9 | 10.3 | 5.5 | 5.4 | 3.6 | 9.4 | 6.4 | 7.9 | 11.6 | 13.5 | 14.1 |
| Lpin1 | NM_001130412 | 0.05 | 2.29E-04 | 24 | 15 | 4.3 | 3.8 | 3.6 | 2.7 | 1.9 | 7.2 | 7.3 | 66.9 | 50.2 | 17.8 | 15.8 | 5.1 | 4.0 | 3.4 | 2.8 |
| Ubqln1 | NM_152234 | 0.05 | 2.29E-04 | 24 | 0 | 3.9 | 29.4 | 29.1 | 21.9 | 19.8 | 16.6 | 14.5 | 14.1 | 12.6 | 28.3 | 18.5 | 21.0 | 21.0 | 25.9 | 23.5 |
| Nmrk1 | NM_145497 | 0.05 | 2.29E-04 | 24 | 18 | 3.9 | 6.0 | 5.5 | 3.5 | 4.3 | 3.9 | 3.4 | 14.9 | 9.5 | 16.2 | 11.6 | 7.0 | 7.2 | 4.5 | 5.5 |

| | | | | | | | | | | | | | | | | | | | | |
|---|---|---|---|---|---|---|---|---|---|---|---|---|---|---|---|---|---|---|---|---|
| Anp32a | NM_009672 | 0.05 | 2.29E-04 | 24 | 0 | 3.4 | 39.8 | 39.6 | 29.9 | 21.8 | 20.0 | 17.0 | 19.8 | 15.4 | 21.0 | 20.0 | 17.3 | 31.3 | 37.0 | 32.9 |
| Ppm1b | NM_001159496 | 0.05 | 2.29E-04 | 24 | 0 | 2.9 | 16.6 | 16.0 | 13.9 | 9.3 | 10.3 | 7.2 | 8.3 | 5.6 | 12.5 | 8.9 | 11.1 | 11.7 | 15.7 | 14.5 |
| Tnfaip2 | NM_009396 | 0.05 | 2.29E-04 | 24 | 1 | 2.8 | 10.6 | 8.3 | 7.3 | 6.6 | 4.0 | 2.8 | 2.5 | 1.4 | 4.3 | 2.6 | 4.7 | 5.7 | 8.0 | 6.4 |
| Tlcd1 | NM_026708 | 0.05 | 2.29E-04 | 24 | 18 | 2.7 | 7.5 | 4.4 | 4.9 | 3.9 | 6.0 | 5.0 | 8.1 | 6.4 | 10.6 | 8.7 | 9.0 | 8.4 | 7.1 | 6.2 |
| Pabpn1 | NM_019402 | 0.05 | 2.29E-04 | 24 | 0 | 2.6 | 15.8 | 14.0 | 10.4 | 8.2 | 7.5 | 6.8 | 5.5 | 5.6 | 12.6 | 7.6 | 9.5 | 13.2 | 13.2 | 8.6 |
| Tspan33 | NM_146173 | 0.05 | 2.29E-04 | 24 | 0 | 2.3 | 9.3 | 8.4 | 7.2 | 5.7 | 4.6 | 3.7 | 3.6 | 3.1 | 5.6 | 3.5 | 6.7 | 5.0 | 7.0 | 8.0 |
| Acnat1 | NM_001164565 | 0.05 | 2.29E-04 | 24 | 0 | 2.1 | 11.9 | 10.6 | 7.1 | 6.3 | 4.9 | 3.7 | 3.3 | 2.3 | 7.8 | 6.8 | 4.9 | 7.9 | 10.9 | 8.0 |
| 40057 | NM_001113487 | 0.05 | 2.29E-04 | 24 | 0 | 2.1 | 11.8 | 10.4 | 7.7 | 6.8 | 5.2 | 4.3 | 3.3 | 2.3 | 7.9 | 6.4 | 8.1 | 7.7 | 9.1 | 7.3 |
| Lpin2 | NM_022882 | 0.05 | 2.29E-04 | 24 | 14 | 2.1 | 4.4 | 4.9 | 3.6 | 3.8 | 6.8 | 5.6 | 10.9 | 8.6 | 8.9 | 7.8 | 4.9 | 5.4 | 4.0 | 2.9 |
| Mtap | NM_024433 | 0.05 | 2.29E-04 | 24 | 0 | 2.0 | 16.0 | 14.0 | 7.4 | 7.6 | 6.5 | 6.1 | 5.6 | 4.8 | 11.2 | 6.1 | 7.8 | 10.2 | 10.1 | 13.9 |
| Taf15 | NM_027427 | 0.05 | 2.29E-04 | 24 | 0 | 1.7 | 17.6 | 14.4 | 8.6 | 6.8 | 6.1 | 7.2 | 8.0 | 5.6 | 9.2 | 8.6 | 9.6 | 10.4 | 13.6 | 9.6 |
| Rpa1 | NM_001164223 | 0.05 | 2.29E-04 | 24 | 0 | 1.5 | 11.6 | 10.2 | 8.1 | 6.3 | 5.0 | 6.2 | 5.4 | 4.6 | 6.0 | 6.9 | 6.0 | 7.7 | 9.5 | 8.3 |
| Lnx2 | NM_080795 | 0.05 | 2.29E-04 | 24 | 0 | 1.4 | 7.3 | 6.8 | 3.8 | 3.7 | 2.1 | 2.1 | 1.3 | 1.5 | 6.5 | 3.2 | 4.1 | 3.4 | 5.9 | 4.3 |
| Gca | NM_145523 | 0.05 | 2.29E-04 | 24 | 0 | 1.2 | 3.6 | 5.4 | 2.9 | 2.5 | 1.3 | 1.5 | 0.1 | 1.2 | 2.3 | 2.3 | 3.1 | 2.5 | 2.0 | 4.0 |
| Bco2 | NM_133217 | 0.05 | 2.29E-04 | 24 | 0 | 1.1 | 6.2 | 6.3 | 4.2 | 3.3 | 4.1 | 2.7 | 1.9 | 1.7 | 3.0 | 2.8 | 2.9 | 4.6 | 5.8 | 4.5 |
| Itgb5 | NM_001145884 | 0.05 | 2.29E-04 | 24 | 0 | 0.9 | 8.4 | 6.8 | 4.1 | 4.5 | 3.4 | 2.8 | 3.2 | 2.2 | 4.8 | 3.4 | 3.5 | 4.8 | 6.4 | 5.2 |
| Sorbs2 | NM_001205219 | 0.05 | 2.29E-04 | 24 | 0 | 0.9 | 5.1 | 4.3 | 3.9 | 3.9 | 3.0 | 3.2 | 2.0 | 1.7 | 4.6 | 3.3 | 4.1 | 4.8 | 5.0 | 4.2 |
| Arntl | NM_007489 | 0.05 | 2.29E-04 | 24 | 0 | 0.9 | 2.3 | 2.9 | 2.1 | 1.2 | 0.2 | 0.2 | 0.3 | 0.0 | 0.7 | 0.8 | 1.0 | 1.5 | 2.1 | 1.0 |
| 40787 | NM_001009818 | 0.05 | 2.29E-04 | 24 | 0 | 0.7 | 5.0 | 4.3 | 3.2 | 2.3 | 1.9 | 2.2 | 1.6 | 1.1 | 2.4 | 4.4 | 3.2 | 3.3 | 3.4 | 4.0 |
| Shkbp1 | NM_138676 | 0.05 | 2.29E-04 | 24 | 0 | 0.7 | 2.7 | 2.3 | 1.7 | 1.5 | 1.1 | 0.9 | 1.0 | 0.9 | 1.7 | 1.1 | 2.1 | 1.6 | 2.0 | 1.7 |
| Syne1 | NM_001079686 | 0.05 | 2.29E-04 | 24 | 4 | 0.5 | 1.4 | 1.8 | 2.1 | 1.5 | 1.1 | 0.8 | 0.6 | 0.3 | 0.6 | 0.5 | 0.6 | 1.0 | 1.8 | 1.7 |
| Rad9a | NM_011237 | 0.05 | 2.29E-04 | 24 | 22 | 0.5 | 1.4 | 1.1 | 0.6 | 0.6 | 0.1 | 0.2 | 0.3 | 0.3 | 0.8 | 0.7 | 1.0 | 1.0 | 1.3 | 0.7 |
| Smarcal1 | NM_018817 | 0.05 | 2.29E-04 | 24 | 0 | 0.4 | 3.5 | 3.3 | 2.3 | 2.1 | 1.7 | 1.5 | 0.9 | 1.0 | 2.0 | 2.1 | 1.9 | 2.1 | 2.8 | 2.3 |
| Gtf3c5 | NM_148928 | 0.05 | 2.29E-04 | 24 | 0 | 0.4 | 3.9 | 2.5 | 1.8 | 1.8 | 1.4 | 1.4 | 0.7 | 1.1 | 1.6 | 1.4 | 1.4 | 2.2 | 2.4 | 1.7 |
| Ptpn9 | NM_019651 | 0.05 | 2.29E-04 | 24 | 0 | 0.3 | 3.3 | 2.3 | 1.3 | 1.6 | 0.8 | 1.1 | 1.0 | 1.1 | 1.9 | 1.1 | 1.3 | 2.3 | 2.1 | 2.9 |
| Ddhd1 | NM_001039106 | 0.05 | 2.29E-04 | 24 | 0 | 0.3 | 2.1 | 1.9 | 0.9 | 1.0 | 0.6 | 0.6 | 0.8 | 0.5 | 1.4 | 0.8 | 0.9 | 1.1 | 1.5 | 1.4 |
| E4f1 | NM_007893 | 0.05 | 2.29E-04 | 24 | 0 | 0.3 | 1.3 | 0.9 | 0.6 | 0.6 | 0.4 | 0.5 | 0.4 | 0.3 | 0.8 | 0.5 | 1.0 | 0.8 | 1.1 | 0.7 |
| Sorbs2 | NM_172752 | 0.05 | 2.29E-04 | 24 | 0 | 0.2 | 2.2 | 2.6 | 1.8 | 1.4 | 1.4 | 1.6 | 0.9 | 0.9 | 1.7 | 1.5 | 1.5 | 2.0 | 2.0 | 2.1 |
| Opn3 | NM_010098 | 0.05 | 2.29E-04 | 24 | 0 | 0.1 | 0.8 | 0.6 | 0.1 | 0.2 | 0.1 | 0.2 | 0.0 | 0.0 | 0.3 | 0.1 | 0.2 | 0.3 | 0.4 | 0.5 |

**Table S3 | Sequence statistics for ribosome profiling data for wild-type (M1W, M3W, M6W) and *Per2* uORF mutant (M15, M20D, M21D) showing the number of reads per sample, number of reads mapped to rRNA, unmapped reads, and mapped reads.**

| Sample | Reads | rRNA aligned | rRNA (%) | Unmapped | Unmapped (%) | Mapped | Mapped (%) |
|---|---|---|---|---|---|---|---|
| M1W | 31067707 | 13722226 | 44 | 3495502 | 11 | 13849979 | 45 |
| M3W | 30481515 | 12873480 | 42 | 3235911 | 11 | 14372124 | 47 |
| M6W | 37574250 | 18179097 | 48 | 3539752 | 9 | 15855401 | 42 |
| M15D | 38245244 | 18128177 | 47 | 3624572 | 9 | 16492495 | 43 |
| M20D | 42932671 | 19864776 | 46 | 4212170 | 10 | 18855725 | 44 |
| M21D | 33744849 | 14525219 | 43 | 3601489 | 11 | 15618141 | 46 |

**Dataset S1 (separate file).** JTK analysis of ribosome profiling data for all transcripts.

**Dataset S2 (separate file).** JTK analysis of ribosome profiling data for all uORFs.

**Dataset S3 (separate file).** Translation efficiency using RNA-seq data from (13) and ribosome profiling data from this study.

**Dataset S4 (separate file).** Ribosome profiling, total RNA sequencing, and translation efficiency from the livers of three wild-type (M1W, M3W, and M6W) and three *Per2* uORF mutant (M15D, M20D, and M21D) mice sacrificed at ZT02-04.

**Dataset S5 (separate file).** Differential expression analysis of total RNA from the livers of three wild-type (M1W, M3W, and M6W) and three *Per2* uORF mutant (M15D, M20D, and M21D) mice sacrificed at ZT02-04.
